# An alpaca-derived nanobody recognizes a unique conserved epitope and retains potent activity against the SARS-CoV-2 omicron variant

**DOI:** 10.1101/2022.12.27.521990

**Authors:** Naphak Modhiran, Simon Malte Lauer, Alberto A Amarilla, Peter Hewins, Sara Irene Lopes van den Broek, Yu Shang Low, Nazia Thakur, Benjamin Liang, Guillermo Valenzuela Nieto, James Jung, Devina Paramitha, Ariel Isaacs, Julian de Sng, David Song, Jesper Tranekjær Jørgensen, Yorka Cheuquemilla, Jörg Bürger, Ida Vang Andersen, Johanna Himelreichs, Ronald Jara, Ronan MacLoughlin, Zaray Miranda-Chacon, Pedro Chana-Cuevas, Vasko Kramer, Christian M.T. Spahn, Thorsten Mielke, Alexander A Khromykh, Trent Munro, Martina Jones, Paul R Young, Keith Chappell, Dalan Bailey, Andreas Kjaer, Matthias Manfred Herth, Kellie Ann Jurado, David Schwefel, Alejandro Rojas-Fernandez, Daniel Watterson

## Abstract

The SARS-CoV2 Omicron variant sub-lineages spread rapidly through the world, mostly due to their immune-evasive properties. This has put a significant part of the population at risk for severe disease and underscores the need for anti-SARS-CoV-2 agents that are effective against emergent strains in vulnerable patients. Camelid nanobodies are attractive therapeutic candidates due to their high stability, ease of large-scale production and potential for delivery via inhalation. Here, we characterize the RBD-specific nanobody W25, which we previously isolated from an alpaca, and show superior neutralization activity towards Omicron lineage BA.1 in comparison to all other SARS-CoV2 variants. Structure analysis of W25 in complex with the SARS-CoV2 spike surface glycoprotein shows that W25 engages an RBD epitope not covered by any of the antibodies previously approved for emergency use. Furthermore, we show that W25 also binds the spike protein from the emerging, more infectious Omicron BA.2 lineage with picomolar affinity. *In vivo* evaluation of W25 prophylactic and therapeutic treatments across multiple SARS-CoV-2 variant infection models, together with W25 biodistribution analysis in mice, demonstrates favorable pre-clinical properties. Together, these data endorse prioritization of W25 for further clinical development.

## INTRODUCTION

Since emergence in late 2019, the highly transmissible coronavirus—severe acute respiratory syndrome coronavirus 2 (SARS-CoV-2)—has infected more than 300 million individuals and claimed more than 6.5 million lives (WHO, updated September 2022). Despite the rapid development of vaccine strategies and preventive measures that help curb disease progression, viral transmission persists, and viral variants with altered virulence and antigenic sites have emerged in successive waves across the globe. Understanding the antigenic nature of emerging variants and the development of broad-spectrum therapeutics and vaccines remains an urgent priority.

SARS-CoV-2 expresses a surface spike (S) glycoprotein, which consists of S1 and S2 subunits that form a homotrimeric viral spike which serves as both viral attachment protein and membrane fusogen ^1^. Cell receptor interaction is mediated by the S1 receptor binding domain (RBD), which binds the peptidase domain of human angiotensin-converting enzyme 2 (hACE2). Structural studies have revealed different conformations of the spike from both in vitro and in situ ^2–4^. In the prefusion stage, the RBD switches between a closed and an open conformation to facilitate hACE2 interaction ^5^. Subsequently, the S2 undergoes a substantial conformational change which drives the fusion of viral and cellular membranes ^1^.

Investigations of sera from COVID-19 convalescent patients enabled isolation of potent neutralizing antibodies (NAbs), the majority of which target the RBD ^6–13^. Some of these have received Emergency Use Authorization (EUA) from the U.S. Food & Drug Administration for SARS-CoV2 treatment or pre-exposure prophylaxis and have shown to be effective in treating certain patients with COVID-19 (https://ww.fda.gov). However, virus evolution causes ongoing emergence of novel SARS-CoV2 variants of concern (VoC), concomitant with reduction of vaccine ^14,15^ and therapeutic antibody efficacy ^16–18^.

The latest SARS-CoV-2 variant of concern (VOC) B.1.1.529, designated Omicron (O), was first reported to WHO from South Africa at the end of November 2021 and is rapidly spreading across many countries, thereby replacing the already highly transmissible SARS-CoV-2 Delta variant (B.1.617.2) ^19^. SARS-CoV-2 Omicron accumulated a large number of mutations compared to its earlier pandemic variants, of which >30 substitutions, deletions or insertions are located in the spike protein. Scientists all over the world are unanimously reporting significantly reduced efficacies of vaccine-elicited sera against Omicron ^20–22^. To best of our knowledge, the majority of potent monoclonal NAbs, including EUA NAbs, showed strongly reduced or no detectable neutralization activity towards Omicron. A notable exception is EUA NAb sotrovimab (S309), which however still exhibits 2-3-fold reduced neutralizing activity ^23^. Accordingly, the identification of novel, potent anti-Omicron NAbs would be highly desirable to strengthen the therapeutic repertoire.

Nanobodies are considered the smallest antigen-binding entity, representing the variable heavy chain fragment (V_H_H) derived from heavy-chain-only antibodies found in camelids (llamas, alpacas, guanacos, vicuñas, dromedary and camels). Because of their small size (2.5 nm by 4 nm; 12–15 kDa) and unique binding domains, nanobodies offer many advantages over conventional antibodies including the ability to bind cryptic epitopes not accessible to bulky conventional antibodies, high tissue permeability, ease of production, and thermostability. Due to their superior stability, nanobodies are highly suited for development as potential bio-inhaled therapies against respiratory diseases. ALX-0171, an inhaled anti–respiratory syncytial virus (RSV) nanobody, demonstrated robust antiviral effect, reduced symptoms of virus infection in animal models ^24,25^ and displayed promising result in reducing RSV viral load ^26^.

We recently described the neutralizing RBD-specific nanobody ^27^, W25, derived from an alpaca immunized with the S protein from the ancestral SARS-CoV-2 strain (Wuhan, Wu). Here, we extend these early findings and show that unlike all hitherto described NAbs, W25 neutralizes the Omicron variant even more potently than the ancestral isolate. In order to understand the molecular basis of neutralization activity and breadth of W25, we determined the cryo-EM structures of W25 in complex with SARS-CoV-2 spike from both ancestral and Omicron SARS-CoV-2, revealing that the vast majority of mutations in the Omicron variant RBD are outside of the W25-RBD binding interface. In addition, W25-binding and live virus neutralization assays demonstrated activity against a broad range of SARS-CoV-2 VOCs. Functional assays reveal that W25 triggers fusion, and that this mechanism is conserved across VOCs, including the less fusogenic Omicron variant. Furthermore, we demonstrate a protection and robust reduction of viral burden and prevention of lung pathology in the K18-human ACE2 mouse model of SARS-CoV-2 challenged with VOC Beta from both prophylactic and therapeutic administration of human IgG Fc-nanobody fusion, W25-Fc. Importantly, W25-Fc treatment results in survival of K18-human ACE2 mouse challenged with lethal dose of Beta SARS-CoV-2. Finally, we analyzed the pharmacodynamics of radiolabelled W25-Fc in mice to gain a better understanding of the pharmacological characteristics of W25-Fc to inform further drug development efforts.

## RESULTS

### W25 ultra-potently neutralizes the SARS-CoV2 Omicron variant

We have previously reported the neutralization activity of W25-Fc against SARS-CoV-2 Wu and D614G ^27^. Given the number of substitutions in the Omicron BA.1 spike protein, including G339D, S371L, S373P, S375F, K417N, N440K, G446S, S477N, T478K, E484A, Q493R, G496S, Q498R, N501Y and Y505H and BA.2 spike including G339D, S371F, S373P, S375F, T376A, D405N, R408S, K417N, N440K, S477N, T478K, E484A, Q493R, N501Y and Y505H in the receptor binding domain, neutralisation activity of W25-Fc was tested against SARS-CoV-2 Omicron BA.1 and BA.2 live virus on Vero E6 cells ^28^. Notably, W25-Fc completely and potently neutralised the Omicron BA.1 and BA.2 variant, with an IC_50_ of 1.45 nM ± 0.31 nM and 2.07 nM ± 0.66 nM, respectively which is ~7 fold more potent compared to the ancestral strain (Fig.1A, B and C). In comparison, EUA mAbs including REGN10933, REGN10987, LY-CoV555 and CT-P59 significantly or completely lost neutralising activity against Omicron BA.1 and BA.2. Other RBD-specific mAbs including CB6 and DH1047 displayed a significant reduction of neutralising activity against Omicron BA.1 variant, by ~4- and 7-fold compared to the Wu strain, respectively (Fig. 1a, b and c) and DH1047 completely lost neutralizing activity against BA.2. Interestingly, the neutralization profiles demonstrated that while W25-Fc exhibited complete inhibition of Wu-1 and Omicron infection, S309, a broadly neutralizing mAb, did not reach 100% neutralization, as seen previously ^29^ (Fig. 1A).

**Fig. 1.**
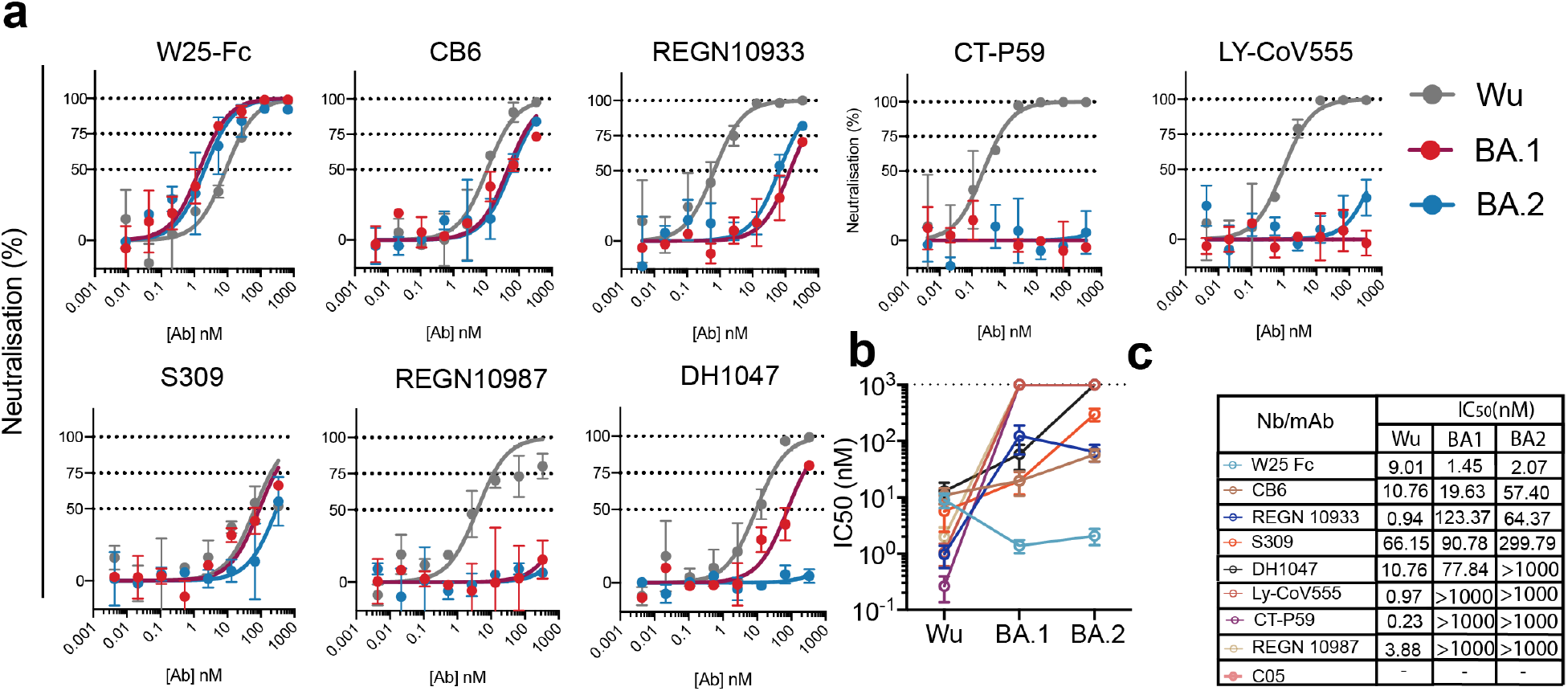
Authentic virus neutralization of SARS-CoV-2 variants by W25, EUA and other mAbs. **a** Neutralization curves comparing the sensitivity of SARS-CoV-2 strains Wu (grey line) and Omicron BA.1 (red line) and Omicron BA.2 (blue line) to antibody as indicated including W25-Fc, C05, CB6, REGN10933, CT-P59, LY-CoV555, S309, REGN10987 and DH1047. The data were analysed and plotted using nonlinear regression (curve fit, three parameters). **b and c** Comparison of IC_50_ values across mAbs tested between Wu and Omicron. IC_50_ was calculated from the neutralization curve by Graphpad Prism 8 software. Color represents different mAbs as indicated. Data are generated from two independent experiments, each performed in technical triplicate.

### Structural analysis of W25 binding to SARS-CoV-2 spike

To gain mechanistic understanding of how W25 binds SARS-CoV-2 Wu and Omicron spikes, we determined their cryo-EM structures in complex with W25 (Table S1). After extensive 2D and 3D classification, two major conformational states of the Wu spike/W25 complex were identified (Fig. S1). Class 1 spike trimers displayed two receptor-binding domains (RBD) in an upward-facing position, with the third one in “down”-position, while class 2 trimers possessed one “up”-RBD and two “down”-RBDs (Fig. S1). For both classes, cryo-EM density maps showed additional density adjacent to the “up”-RBDs, not covered by the structural model of the spike trimer, strongly suggesting that these density portions correspond to W25 (Fig. 2a, upper panel, and Fig.S2). For the Omicron spike/W25 complex, multiple rounds of 2D and 3D classification identified “3 down”, “2 up, 1 down” and “1 up, 2 down” RBD classes (Fig. S3). The latter state displayed clear extra density corresponding to W25 in similar position as for the Wu spike (Fig. 2a, lower panel).

**Fig. 2.**
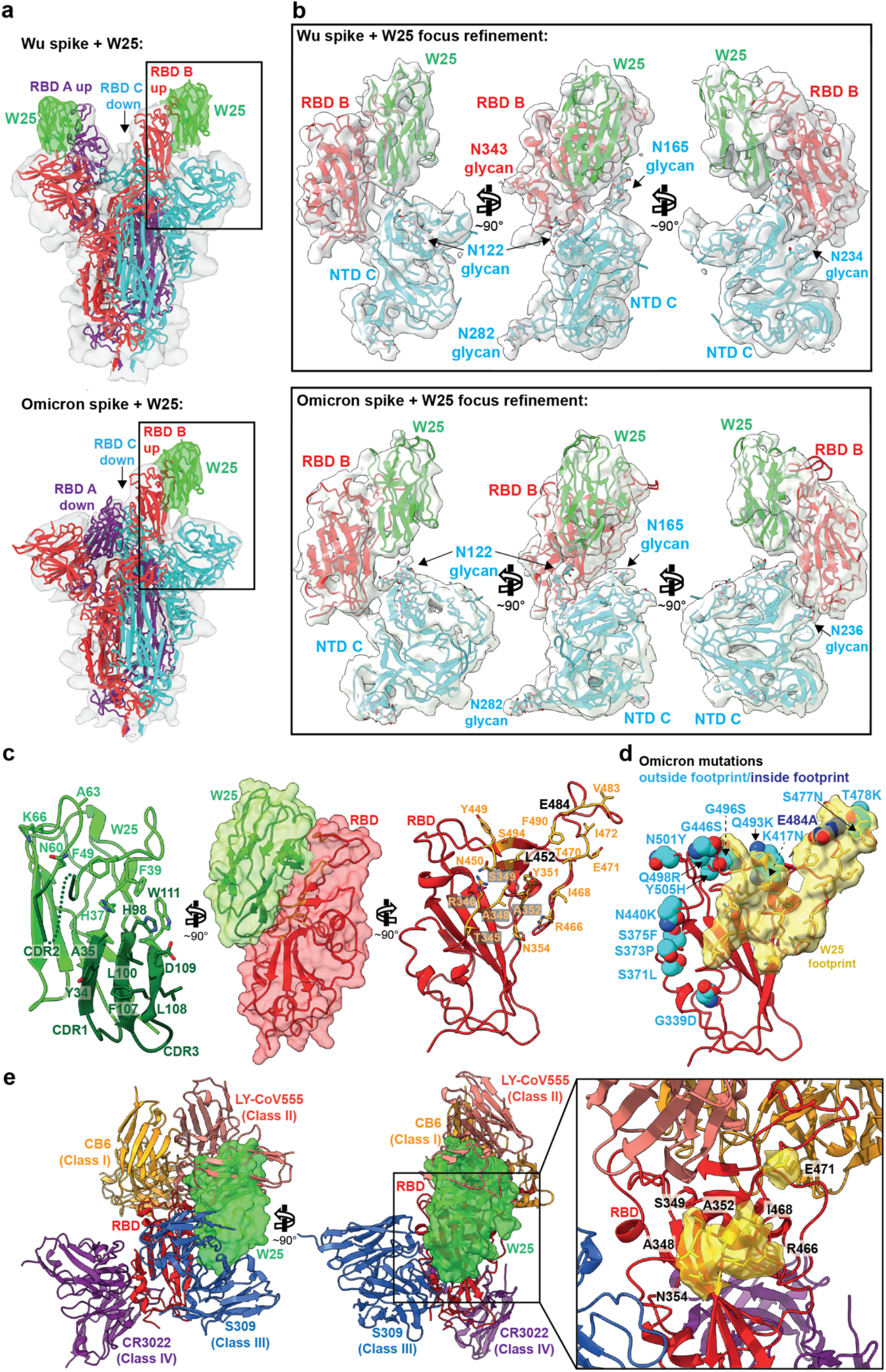
Structure analysis of the W25/spike interaction. **a** Cryo-EM densities of Wu (upper panel) and Omicron (lower panel) spike/W25 complexes (grey semi-transparent surfaces) (map 1, Fig. S1 and map 4, Fig. S3). Fitted trimeric SARS-COV2 spike protein structures (PDB 6ZXN (Wu) and 7WG6 (Omicron) ^63,78^ are shown in cartoon representation, with protomer A colored purple, protomer B red and protomer C cyan. W25 is shown as green cartoon. Densities corresponding to W25 are colored green. **b** Three views of a focused refinement of the regions illustrated in A. Proteins are shown in cartoon representation, and N-linked glycans as sticks, in the same colors as in A. Cryo-EM densities (map 3, fig. S1, and map 5, fig. S4) are shown as semi-transparent light gray surfaces. **c** Detailed “open book” view of the W25/RBD interaction interface between RBD and W25. Amino acids involved in intermolecular interaction are shown as sticks. **d** Visualization of the RBD surface in contact with W25 (yellow). The RBD is shown as red cartoon. Residues mutated in the Omicron RBD are shown as spheres, and colored light blue when located outside of the W25 contact surface, or dark blue when inside. **e** Two views of a superposition of selected RBD-bound nABs with the RBD/W25 complex, in cartoon representation (CB6 – PDB 7C01, LY-CoV555 – PDB 7l3N, S309 – PDB 7R6W, CR3022 – PDB 6YLA). W25 is shown as semi-transparent green molecular surface. For clarity, only the W25-bound RBD is shown as red cartoon. The inset shows an enlarged view of the unique portion of the W25 epitope as yellow surface.

Focused refinement of the Wu and Omicron spike RBDs, the NTD regions of the adjoining spike protomer, and W25 resulted in contiguous density maps of sufficient quality for flexible fitting and refinement of a W25 structure model generated by AlphaFold2 ^30,31^ (Fig. 2b, Table S1 and movie S1). W25 binds to the side and upper edge of the RBD, with the downward-facing side of W25 surrounded by N-linked glycans arising from spike protomer B NTD N122 and N165 (Fig. 2b). Since W25 positions relative to Wu or Omicron RBDs were equivalent, and the map quality and resolution of the Wu spike/W25 complex was slightly better (Fig. 2b and Fig. S1 and 3), the Wu spike/W25 model was subjected to further structural analysis. Superposition of the RBD/W25 complex structure with the Wu spike class 1 “down”-RBD indicates that W25 binding would lead to a slight overlap with the adjacent spike NTD, rationalizing why W25 preferentially binds to RBDs in the “up”-conformation (Fig. S4A).

Detailed inspection of the RBD/W25 contact surface reveals an interface area of 847 Å^2^ with mostly hydrophobic character, involving W25 β-sheet 3 CDR1 residues Y34, A35 and downstream amino acids H37, F39 (Fig. 2c). In addition, there are interfacing residues in the loop region connecting β-sheets 3 and 4, which include CDR2 (M41, R47, F49, F53, G54, N60, A62, Y61, A63, K66), and ultimately residues in β-sheet 7 (CDR3): H98, L100, L105-W111. From the RBD side, amino acid residues in an N-terminal loop region contribute to W25 binding (T345-S349, Y351, A352, N354), as well as from the loop downstream of β-sheet 3 (Y449-L452) and from a large loop preceding β-sheet 4 (R466, I468, T470-I472, G482-E484, F490, S494). In addition, there are hydrogen bonds between W25 side chains D109, W111 and RBD side chain S349 and the backbone carbonyl group of N450, respectively, as well as between RBD side chains N354, T470 and backbone carbonyls of W25 F53, L105. Twenty-two out of 23 epitope residues are conserved between Wu and Omicron. Only one interfacing residue is non-conservatively exchanged between Wu and Omicron SARS-CoV2 (E484A) (Fig. 2d, Fig. S4b and movie 1.).

### Comparison with RBD binding modes of other nanobodies, neutralizing antibodies and ACE2

In order to put the present structure analysis in context, PDB entries containing camelid or synthetic SARS-CoV2 specific nanobodies were collated and structurally aligned on their RBDs, together with the RBD/W25 complex (Fig. S4c and Table S2). The vast majority of nanobodies either attach to one side of the RBD (“side-1” or cluster 1 in ^32^) or to the upper RBD surface (“top” or cluster 2). Two of the nanobodies bind to an RBD region opposite of cluster 1 (“side-2”). Interestingly, the W25 binding mode seems to incorporate characteristics of two of these clusters by occupying a surface area overlapping with epitopes of both “top” and “side-2” binders.

In addition, structures of the variable fragments of selected nABs, representing distinct RBD-binding modes (classes 1-4) ^33^, were superimposed on the W25/RBD complex structure (Fig. 2e). W25 partially overlaps with the VH fragment of class 2 NAb LY-CoV555, resulting in partially shared epitopes involving RBD amino acid residues 351, 449-453, and 480-490. Furthermore, W25 minimally clashes with loop regions of class 3 NAb S309 VH (residues 102-109) and VL (residue 94); their respective epitopes are adjacent but do not overlap. Importantly, a significant portion of the W25 epitope, including surface-exposed RBD amino acid stretches 348-354 and 466-471, is unique and not covered by the CDRs of the representative class 1-4 NAbs analyzed here (Fig. 2e, right panel).

Subsequently, structural alignment of the RBD moieties of the RBD/W25 complex presented here with the structure of the RBD bound to its human cell surface receptor ACE2 was performed. The analysis demonstrates that there is a slight overlap of the molecular surfaces (Fig. S4d). In this hypothetical model, W25 side chain E46 directly opposes ACE2 residue E35, which might induce electrostatic repulsion and lead to obstruction of the ACE2-binding site or ACE2 displacement from the RBD (Fig. S4e).

### W25 binds multiple VOCs of SARS-CoV-2 and retains binding to BA.2 Omicron sublineage

To evaluate W25 binding to previous SARS-CoV-2 VOCs and the rapidly expanding BA.2 Omicron sublineage, we expressed and purified Wu and VOC spikes (bearing mutations indicated in Fig. S5a), and validated spike purity and trimer formation by SDS-PAGE (Fig. S5b) and SEC (Fig. S5c). Binding kinetics of W25-Fc to the spike variant proteins were assessed by surface plasmon resonance (SPR) (Fig. 3a). Subnanomolar dissociation constants (K_D_) were observed for Wu, Alpha, Beta, Gamma, and Omicron BA.1 using a 1:1 kinetic model (Fig. 3b). The K_D_ values of W25-Fc for Delta and Lambda variants were higher compared to other variants, at 11.4 and 0.7 nM, respectively (Fig. 3a,b). As control, S309 and DH1047 NABs also yielded subnanomolar K_D_ values (Fig. 3b, and Fig. S6). ELISA was consistent with these findings (Fig. 3c and Fig. S7a), showing that W25 binds all variants with similar K_D_ (0.033-0.048 nM), except for the Delta variant (1.8 nM). In agreement with recent reports ^34,35^, and our neutralisation assays, EUA RBD-specific mAbs including REGN10933, LY-CoV555 and CT-P59 showed significantly reduced binding affinity (>10-fold reduction of K_D_) for Omicron BA.1, with the exception of S309 (Fig. 3c). We observed no binding of REGN10987 to Omicron BA.1 variant. Structurally defined cross-reactive mAbs including DH1047 (RBD-specific NAb), CR3022 and 2M10B11 (non-neutralizing cryptic RBD epitopes) showed significantly reduced binding to Omicron BA.1 variant (~5 fold).

**Fig. 3.**
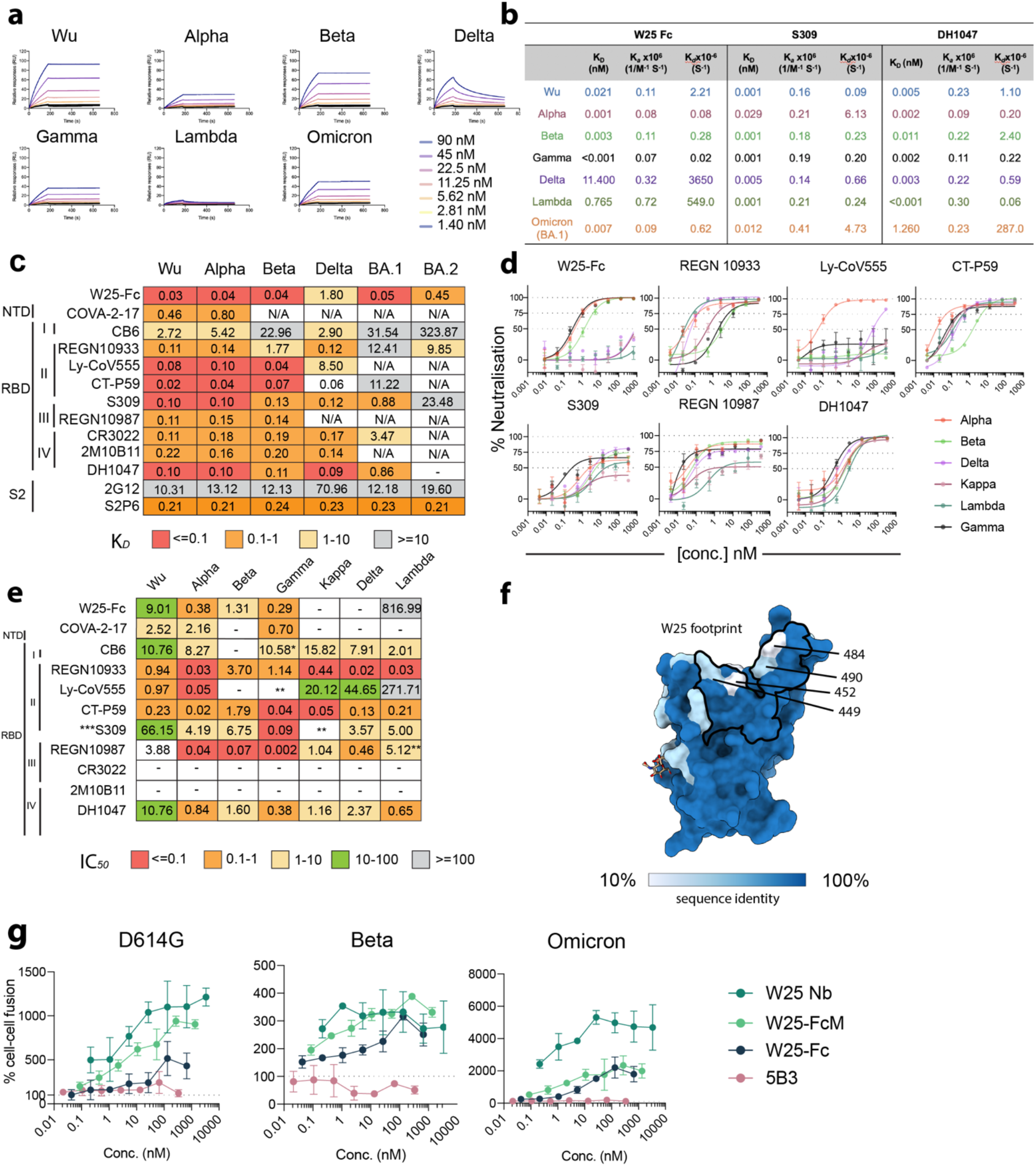
Broad reactivity of W25-Fc is conferred by conserved patch on SARS-CoV-2 RBD and potential inhibition mechanism of W25. **a** SPR sensograms of W25-Fc. W25-Fc was immobilized on SPR protein A chips. Various concentrations of SARS-CoV-2 spike variant proteins as indicated were injected for 30 s, at 30 μL/min followed by dissociation for 600 s. Dissociation constants (*K_D_*) were determined on the basis of fits, applying a 1:1 interaction model. Similar experiments were conducted for S309, DH1047 and C05 (as control, fig. S6). **b** Summary table showing *K_D_, K_a_, K_d_* of indicated Nb/mAbs. **c** K_d_ values from ELISA binding curves of W25-Fc, EUA mAbs other epitope specific mAbs to SARS-CoV-2 Spikes variants. **d** Neutralisation curves comparing the sensitivity of live SARS-CoV-2 viruses including Alpha, Beta, Delta, Kappa, Lambda and Gamma to W25-Fc, EUA and other RBD-specific mAbs as indicated. The data were analysed and plotted using nonlinear regression (curve fit, three parameters) and IC_50_ value was calculated from neutralisation curves by Graphpad Prism 8 software. **e** Summary of IC_50_ values (nM) of neutralistion of SARS-CoV-2 variants performed in VeroE6 cells. Values that approached 50% neutralization were estimated from (D). Data are generated from two independent experiments, each performed in technical duplicate. **f** Molecular surface conservation of RBD VOCs. **g** W25 and its derivatives enhance spike-mediated cell fusion. Cell-cell fusion assay was performed. Stable cells expressing the rLuc-GFP components (effector cells) were transiently transfected SARS-CoV-2 spike proteins bearing spike mutations for D614G (top left), Beta (top right), Omicron (Bottom left) or control plasmid (no envelope control). W25 Nb, W25 Dimeric Fc (W25-Fc) or W25 monomeric Fc (W25-FcM) were incubated with effector cells for 1 hour prior to co-culturing with target cells (huACE2-HEK293T cells). After 24-48 hours, *Renilla* luciferase was read and analysed by subtraction with control plasmid treatment. Data are generated from three biological replicates.

Continuing surveillance of Omicron evolution revealed the newly emerged BA.2 Omicron sublineage, containing 8 unique spike mutations while lacking 13 spike mutations found in BA.1 ^29^. Of the anti-RBD panel we tested, only W25-Fc retained sub-nanomolar affinity to BA.2 with a K_D_ value of ~0.4 nM. Conversely, all EUA mAbs, including S309, showed significantly impaired binding to BA.2 spike protein (Fig. 3c and Fig. S7b). S2-specific mAbs S2P6 and 2G12 bound all variants with K_D_’s of approximately 0.2 nM and 10-12 nM, respectively (Fig. 3c). Additionally, a direct comparison with other previously characterised engineered Nb-Fc fusions demonstrated that W25-Fc exhibits superior binding affinity for Omicron BA.1 (Fig. S7c).

### W25 neutralizes multiple VOCs of SARS-CoV-2

As SARS-CoV-2 continues to evolve, the dominant variants are continuously changing. To explore and compare the efficacy of W25 for neutralization of SARS-CoV-2 VOCs, we performed live virus neutralization on Vero E6 cells against major VOCs including Alpha, Beta, Gamma, Delta, Kappa and Lambda and found that W25-Fc neutralized Alpha, Beta and Gamma variants (Fig. 3d, e). W25-Fc showed complete and potent inhibition of SARS-CoV-2 Alpha, Beta, Gamma variant infections with IC_50_ values ranging from 0.38-1.31 nM. The monomeric W25 also neutralised similar VOCs with IC_50_ values ranging from 0.42-2.12 nM (Fig. S8a,b). Interestingly, both monovalent and divalent forms exhibited significantly weaker neutralisation for Kappa, Lambda, and Delta variants. Of note, monomeric W25 was effective against Beta and Gamma, two of the most resistant variants leading to first-generation RBD-associated antibodies ^15^. Similar to other reports, we demonstrated that introducing bivalency through Fc fusion further improved efficacy of W25-Fc (~10 fold for Gamma variant). Overall, the neutralisation data are consistent with the W25 binding mode observed in the cryo-EM analysis, since key residues within the W25 binding interface, with the exception of L452 and E484, were highly conserved across all VOCs examined in this work (Fig. 3f and Fig. S9).

### W25 stimulates spike-dependent cell-cell fusion

We furthermore explored the modulation of spike-mediated membrane fusion as a potential neutralization mechanism, using established cell-cell fusion assays ^36^. W25 and its derivatives (Fc-fusion and monomeric Fc-fusion [FcM]) caused extensive cell-cell fusion for SARS-CoV2 D614G-, Beta- and Omicron-infected cells, demonstrated by luciferase activity (Fig. 3g), when compared to a control antibody (anti-NiV F, 5B3). This was most striking for Omicron, which in the absence of W25 did not cause significant cell-cell fusion. Overall, W25-Fc induced less spike-mediated fusion, followed by W25-Fc monomeric (FcM) and W25 (W25 Nb) in all variants tested. No enhancement was observed for control antibody treatment. Additionally, live cell GFP-reporter IncuCyte assays were performed in parallel and showed consistent results, where W25 addition greatly enhanced the GFP signal (Fig. S10 and 11). Thus, W25 may additionally act through premature fusion activation, causing irreversible inactivation of coronavirus particles.

### W25 prophylactically and therapeutically reduces disease burden and protects from SARS-CoV-2 infection mortality in mice

Since nanobody affinity and function have shown to be limited by temperature ^37^, thermostability of W25 was tested and showed W25 retains its antigen-binding function after heat treatment at 90°C (Fig. S12a,b). In addition, full stability of the monomeric W25 after nebulization was observed, suggesting potential stability for airway administration (Supplementary Fig.12c,d). To assess the use of W25-Fc for pre- and post-exposure therapy to counter SARS-CoV-2 infection, the K18-hACE2 transgenic murine model was employed. W25-Fc or GFP-Nb-Fc (control) were administered by intraperitoneal injection (i.p.) 4 h prior to or 24 h after intranasal (i.n.) challenge of K18-hACE2 mice with SARS-CoV-2 Beta variant (Fig. 4a, f). It has been shown previously that SARS-CoV-2 Beta is significantly more lethal and causes more severe organ damage in K18-hACE2 mice than both the Wu ancestral strain and the D614G variant, mimicking severe SARS-CoV-2 infection in humans ^38^. Infected mice were monitored for 10 or 14 days after infection. All SARS-CoV-2-infected animals from the prophylactical W25-Fc treatment group survived, as opposed to control-treated group (Fig. 4b). Symptoms, including ruffled fur/piloerection and accelerated shallow breathing, were observed in control-treated animals at 4-5 days after infection and progressed rapidly. Some non-treated, infected mice reached the humane endpoint state by day 6 post infection. Prophylactical W25-Fc treatment prevented all respiratory disease (Supplementary Fig.13) and weight loss (Fig. 4d) caused by SARS-CoV-2 Beta infection. In a separate experiment, lungs were collected after 4 days of infection. W25-Fc treatment reduced infectious virus level up to 10^3^-fold when compared to the control group, and viral RNA also showed a significant reduction (Fig. 4d, e). Therapeutic treatment (24 h post infection) of SARS-CoV-2 Beta infected mice (Fig. 4f) with W25-Fc significantly improved survival (60%) (Fig. 4g), weight loss (Fig. 4h) and lead to lower viral load in nasal turbinate (Fig.4i) and lung (Fig. 4i), compared to GFP-Nb-Fc treated infected mice.

**Fig. 4.**
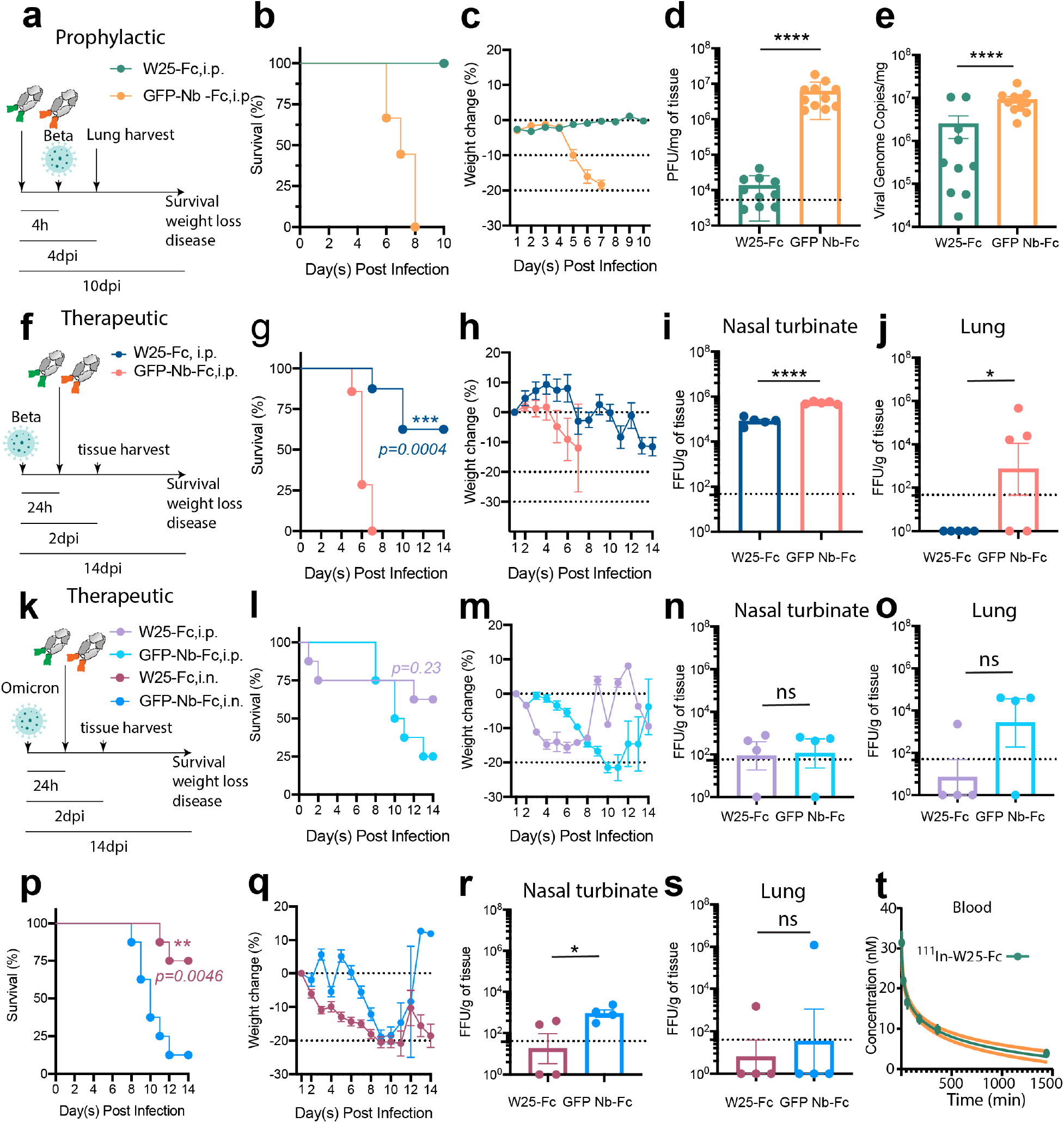
W25-Fc protects mice from lethal Beta and Omicron SARS-CoV-2 infection and W25-Fc pharmacodynamic analysis. **a** Experimental schematic: eight-to twelve-week-old male and female K18 transgenic mice were inoculated via the intranasal route with 1×10^3^ PFU of SARS-CoV-2 (Isolate B.1.351). W25-Fc or GFP Nb-Fc were administered intraperitoneally 4 hours prior to infection. **b** Survival and **c** weight change were monitored and scored. Two experiments were performed (n = 9-12, Log-rank (Mantel-Cox) test *\*\*\*\*p< 0.0001*). **d** Viral burdens were determined in lung tissues 4-days post infection via plaque forming assays for infectious virus and **e** RT-qPCR for viral genome copy number. Two experiments were performed (N = 10–11). **f** Experimental schematic: five to six-week-old female K18 transgenic mice were inoculated via the intranasal route with 1×10^4^ FFU of SARS-CoV-2 (Beta, Isolate B.1.351) N=8, and N=4-5 each group for survival and tissue harvest experiment, respectively. For therapeutic treatment, W25-Fc or GFP-Nb-Fc were administered intraperitoneally 24 hours after SARS-CoV-2 Beta infection at 5×10^3^ FFU/mouse. **g** Survival and **h** weight change were monitored. Viral loaded were determined in **i** nasal turbinate and **j** lung tissues 2-days post infection by immuoplaque assays for infectious virus. **k** Experimental schematic: for Omicron infection, five to six-week-old female K18 transgenic mice were inoculated via the intranasal route with 1×10^5^ FFU of SARS-CoV-2 (Isolate B.1.1.529). W25-Fc or GFP-Nb-Fc were administered intraperitoneally or intranasally 24 hours after infection. **l and p** Survival and **m and q** weight changes were monitored and scored. Viral loaded were determined in **n and r** nasal turbinate and **o and s** lung tissues 2-days post infection by immuoplaque assays for infectious virus. Bars represent medians, dots are individual animals and dotted horizontal lines indicate the limit of detection. **t** The pharmacodynamics of W25-Fc in blood was determined by conjugation of W25-Fc to radioactive ^111^Indium. One mg/kg were injected intravenously through the tail to 6 groups of mice (group 1: 5 min, group 2: 20 min, group 3: 60 min, group 4: 3 hr, group 5: 5 hr and group 6: 24 hr). The mice were dissected and concentration of ^111^In W25-Fc was measured in the blood using an Auto-Gamma Counter.

The efficacy of W25-Fc against Omicron BA.1 was also tested. Omicron virus appears less pathogenic in K18-ACE2 mice ^39^ compared to earlier isolates. In this work, we inoculated mice with a lethal dose (10^5^ FFU/mouse), and W25-Fc was administrated 24 hours after infection via i.p. or i.n. delivery (Fig. 4k). In parallel experiments, tissues were harvested 2 days post infection. For i.p delivery, mice treated with W25-Fc had slight improvement of survival rate as compared to those which received control treatment (62.5% vs 25%,*p*=*0.23*). Some mice treated with W25-Fc reached humane endpoint by 2 days post-treatment (Fig. 4l). Mice-treated with W25 via i.p. displayed weight loss between day 2 to 8 post infection (Fig. 4m). No significant reduction of viral load in nasal turbinate (Fig. 4n) and lung was observed (Fig. 4o). Notably, i.n. delivery of W25-Fc to Omicron-infected mice significantly improved survival (75%) compared to control-treated group (Fig. 4p). Mice that received W25-Fc i.n. had lower weight compared to control treated mice (Fig. 4q), but also lower viremia in nasal turbinate compared to controls (Fig. 4r). No significant difference between W25-Fc and control treated mice was observed for viremia in lung (Fig. 4s). Consistent with other studies, we observed low level of viremia in lung from Omicron-infected mice compared to SARS-CoV-2 Beta variant infection ^39,40^. Finally, we determined the pharmacokinetic profile of single dose intravenously injected W25-Fc radiolabelled with 1.2-1.5 MBq/mouse (1 mg/kg) of indium-111 in C57BL/6 mice (Fig. S14a). *Ex vivo* studies were carried out measuring radioactivity in several tissues including blood, lungs, liver, kidneys, spleen, brain, muscle and tail after 5 minutes, 20 minutes, 1 h, 3 h, 6 h and 24 h (Fig. S14b-h). We found that the half-life of W25-Fc in blood is 20.6 h, and after 24 hours, the concentration of W25-Fc in blood was 3.8 ± 03 nM, well above the *in vitro* neutralization IC_50_ values for most variants (Fig. 3d,E, 4T).

## DISCUSSION

COVID-19 vaccines have proven to be highly effective in reducing the number of severe cases ^41,42^, and are associated with lower transmissibility ^43^, but SARS-CoV-2 continues to evolve. The Delta variant has been dominant around Sep-Dec 2021, but more recently Omicron and its sublineages prevailed due to their unparalleled transmissibility. Certain therapeutic antibody cocktails have been approved for post exposure treatment to reduce severe illness. However, the potency and breadth of protection by the majority of therapeutic antibodies are compromised by emerging SARS-CoV-2 immune escape mutations, underlining an urgent need for developing, preparing and improving antiviral avenues.

The COVID-19 pandemic highlighted the importance of the rapid discovery of antiviral drugs. In this effort, a growing list of neutralising antibodies, nanobodies and synthetic antivirals have been developed using tools such as human/mouse B cells or immunised camelids ^8–11,44^. In contrast to conventional antibodies that require post-translation modifications from mammalian cells, nanobodies can be expressed in prokaryotic cells, hence, are cheaper to produce. Benefiting from their small size, nanobodies may bind with higher affinity, target epitopes not accessible to conventional antibodies, and are amenable to nasal administration to directly reach infected mucosal tissues and lungs ^45^. However, due to their low molecular weight, nanobodies may have very short half-lives in the bloodstream. Our work and previous reports ^46^ showed that dimerization via Fc-fusion significantly improved neutralisation potency, potentially through simultaneous engagement of two adjacent “up”-RBDs (Fig. 3d, e).

The majority of neutralizing antibodies raised against Wuhan SARS-CoV-2 are not effective against SARS-CoV-2 Omicron. In this work, we demonstrate that W25 and W25-Fc display potent inhibitory activity *in vitro* against the SARS-CoV-2 Omicron variant, as well as other VOCs including Alpha, Beta and Gamma (Fig. 1 and 3d, e) in live virus neutralization assays. This is consistent with structure-based sequence alignments (Fig. S9), demonstrating that the majority of W25 residues involved in RBD binding are highly conserved across SARS-CoV-2 VOCs (Fig. 3f), with the exception of L452, which is centrally located in the binding interface. Neutralization of Kappa, Lambda and Delta variants by W25-Fc is strongly impeded (Fig. 3d, e). These variants share the L452R/Q mutation, which may lead to steric clashes with W25 residues F39 and/or W111 (Fig. 2d, 3f and fig. S9), thus rationalizing weaker binding affinity and neutralization activity of W25 towards these variants (Fig. 3c-e). This also suggests that W25 might not be effective against Omicron BA4/5. Unlike other antibodies ^47–51^, mutations at RBD position 484 (E484K/A/Q), presented on Beta, Gamma, Kappa and Omicron spike variants, do not impede the binding and neutralizing activity of W25. These mutations might be tolerated since E484 is only peripherally situated in the W25/RBD interface (Fig. 2d and 3f). Intriguingly, W25, but not S309, retains a relatively high affinity to BA2.1 spike bearing mutation at amino acid 371 which is thought to be the key change due to introduction of a bulky phenylalanine side chain, causing loss of efficacy for several EUA mAbs and nanobodies ^51,52^. This is also in accordance with our structure analysis, showing no involvement of residue 371 in W25 binding. Additionally, recently emerged circulating strains of BA.2 derivatives (BA.2.75 and BA.2.75.2) still retain the W25 key interacting residues, suggesting the neutralizing activity of W25 to these variants is retained.

Several SARS-CoV-2 neutralizing antibodies have been reported targeting NTD and S2 epitopes, the most effective antibody classes however appear to target the RBD due to its natural role in binding the human ACE2 receptor. Prior structural studies have classified these SARS-CoV2 NABs according to their RBD binding modes ^33,53^ (Fig. 2e). Class 1 and 2 NABs obstruct the ACE2 binding site on top of the RBD. However, they are highly susceptible to VOC escape mutations including K417N, G446S, E484A and Q493R. Classes 3 and 4 bind largely outside of the ACE2 interaction interface, to opposite surfaces on the side of the RBD. Our structure analysis showed that W25 targets a unique RBD epitope, partially overlapping with class 2 and 3 NABs, not affected by Omicron mutations. Accordingly, W25 might be suitable as therapeutic agent in combination with class 1 and/or 4 NABs to improve treatment efficacy against SARS-CoV2 Omicron strains or newly emerging VOCs.

Most N-glycosylation sites are highly conserved across multiple SARS-CoV-2 variants, particularly in the NTD (N61, N122, N165 and N234). Our structural analysis found that W25 is surrounded by N-linked glycans at N122 and N165 from NTD of the neighboring spike protomer (Fig. 2b). Interestingly, beyond its shielding function, N165 act as a molecular switch for RBD conformation by helping to maintain RBD in the ‘up’ or ACE2-accessible state ^54,55^. Accordingly, RBD/W25-interaction might interfere with these glycan-mediated RBD opening processes. In addition, the binding of a class 2 NAb, BD-23, is also facilitated by this glycan ^56^.

The mode of action of W25 in activating fusion mechanism may have clinical implications. Our structural analysis indicates that W25 only associates with RBD-up state and slightly overlaps with the ACE2 binding surface. This is in line with our fusion assay demonstrating that W25 triggers fusion activity of transfected Spike protein in the presence of the cognate receptor ACE2 on the target cell population. Thus W25 may promote RBD-up conformation facilitating ACE2 engagement and enhance fusion. However, no infectivity enhancement was observed in authentic virus neutralization assay (Fig. 3d, e), thus W25 may trigger premature fusogenic conformational changes on spike leading to viral inactivation prior to cellular engagement. A similar mechanism has been reported for RBD specific mAbs including antibodies CR3022 ^57^ and S230 ^58^ and nanobodies such as VHH E, W and U ^51^. However, the mode of engagement of W25 to spike is unique to these previously reported binders, and incorporates the outward face of RBD and is largely non-overlapping with the ACE2 site (fig. S15). Interestingly, induction of fusion was reduced using the dimeric W25-Fc relative to the monomeric form, although dimeric W25-Fc showed increased antiviral activity, thus multiple modes of action may underpin the potency of W25.

In K18-hACE2 mouse model, W25-Fc administrated before infection provided full protection to all treated animals from a fatal SARS-CoV-2 challenge dose. Mice that received W25 prophylactic treatment exhibited no sickness and weight loss following the challenge with SARS-CoV-2 Beta variant (Fig. 4a-e). Single-dose of therapeutic treatment of W25 after Beta and Omicron challenge provided improvement in survival (Fig. 4f to s). Interestingly, i.n. delivery of W25 conferred a better outcome compared to that of i.p. delivery This could be attributed to virus tropism where Omicron is reported to have higher replication in *ex vivo* bronchial epithelium as compared to other VOCs and Wu ^59^. This suggests that i.n. delivery of W25 may directly block the key infection site for Omicron. Nebulization, being more efficient at targeting the deep and local pulmonary structures could be tested for W25 efficacy in preclinical setting. W25 in combination with other Nbs could be used as novel cocktail treatment that could be rapidly generated to block virus mutational escape. Of note, the efficiency of delivery methods for each VOC may need to be considered and assessed.

We also recognize that there are some limitations in our present study. Firstly, our experiments analyzing the neutralization profile and cell to cell fusion were performed in VeroE6 cells and in HEK293 cells overexpressing ACE2. While these cells are widely used and validated in preclinical settings, future investigations using primary nasal and/or lung epithelial cells may offer insight into the observed *in vivo* therapeutic activity. Secondly, as nebulization was only performed in uninfected mice, nebulization in an infection model would need to be performed to access the efficacy. Relevant to this, however, i.n. delivery of W25-Fc was shown to be therapeutically beneficial for Omicron infection.

In conclusion, our results show that the W25 nanobody, raised against the RBD of the ancestral SARS-CoV-2 strain, has highly potent neutralizing activity against multiple SARS-CoV-2 variants including Omicron *in vitro*. Our structural analyses reveal that the binding interface of W25 on the RBD is unique and highly conserved across multiple VOCs*. In vivo* mouse model SARS-CoV-2 infection studies demonstrated that W25 confers protection when administrated prophylactically or therapeutically, suggesting its therapeutic potential as a passive immunotherapy. The COVID-19 pandemic is still ongoing, with SARS-CoV-2 constantly evolving, prompting speculation that there will be new VOCs driving a resurgence of infections. Although vaccination remains the main measure against the pandemic, it is not effective in or possible for all patients. Accordingly, there is still an urgent need to provide additional means for infection prevention and/or treatment of high-risk patients and those in mid- to low-income countries. Because of its high efficiency, remarkable stability, and resilience to nebulization W25 has high therapeutic potential in SARS-CoV-2 infected individuals.

## MATERIALS AND METHODS

### Cell lines

Cell lines used in this study were obtained from ATCC (HEK293T and Vero-E6) or Thermo Fisher (ExpiCHO and Flp-In^™^ T-REx^™^ 293). VeroE6 and HEK293T cells were maintained in DMEM supplemented with 10% heat-inactivated (HI) Fetal Calf Serum (FCS, Bovogen), penicillin (100 IU/ml)/streptomycin (100 μg/ml) (P/S, Gibco) and L-glutamine (2 mM) (Life Technologies). Cells were cultured at 37 °C and 5% CO_2_. ExpiCHO cells were maintained in ExpiCHO^™^ Expression media as per the manufacturer’s protocol. ExpiCHO cells were cultured at 37 °C and 7.5% CO_2_. All cell lines used in this study were routinely tested for mycoplasma and found to be mycoplasma-free (MycoAlert Mycoplasma Detection Kit MycoAlert, Lonza). Flp-In^™^ T-REx^™^ 293 cells were cultured according to the manufacturer’s protocol.

### Transient expression of Spike protein, antibodies and nanobodies

#### Expression and purification of SARS-CoV-2 Spike, antibodies and nanobodies

Soluble, trimeric spikes (residue 1-1204 amino acid) of SARS-CoV-2/human/China/Wuhan-Hu-1/2019 (referred as Wu) (Genbank: MN908947), SARS-CoV-2 Beta variant (B.1.351), Alpha variant (B.1.1.7), Gamma variant (P.1), Lambda variant (C.37), Delta variant (B.1.617), Omicron (BA.1) and Omicron (BA.2) spike mutations were added *in silico* into the codon-optimised Wuhan reference stain and were cloned to pNBF plasmid. Spike proteins contain 6 proline mutations (F817P, A892P, A899P, A942P, K986P and V987P) and substituted at the furin cleavage site (residues 682–685) ^60^. For antibodies and nanobodies, DNA encoding antibody variable domains including W25, W270 ^27^, NB3, Nb11, Nb20 ^50^, MR17 ^61^, SB14, SB42, SB4 ^62^, Ty1^63^ and H11 ^64^ and GFP protein were cloned in-frame with an upstream IgG leader sequence and downstream human IgG1 Fc open reading frame into the mammalian expression vector pNBF. For monoclonal antibodies, heavy and light chains of S309 ^9^, REGN10987, REGN10933^10^, Ly-CoV555 ^11^, CT-P59 ^12^, CB6 ^8^, DH1047 ^13^, 5B3 ^65^, and C05 ^66^ were cloned into a human IgG1 expression vector as described previously ^67^. After validation of the cloning by Sanger sequencing, the plasmids were transfected into ExpiCHO cells (Thermo Scientific) according to the manufacturer’s protocol (1 μg DNA/ml of cells; 2:3 ratio of heavy chain to light chain). After 7 days, cell culture media were clarified by centrifugation and the IgG captured using Protein A resin (GE Healthcare). Proteins were eluted from the resin using citrate buffer pH 3, eluate was buffer exchanged to PBS pH7.4. For nanobody production, W25 and W270 were expressed as previously described ^27^. Purified antibodies, and nanobodies were verified by SDS-PAGE Coomassie staining analysis.

### Expression and purification of HexaPro spike protein

A stable Flp-In^™^ T-REx^™^ 293 cell line (Thermo Fisher) expressing the prefusion-stabilized HexaPro variant of the Wu Spike ^68^ was generated according to the manufacturer’s protocol (polyclonal selection). Cells were seeded in T-300 flasks (0.3×10^6^ cells/ml), and after 24 h the medium was exchanged against fresh medium containing 1 μg/ml tetracycline to induce protein expression. After 4 days, cell culture supernatant was dialyzed against binding buffer (50 mM Tris-HCl pH 7.8, 500 mM NaCl), and spike was purified using Strep-Tactin XT beads (IBA Lifesciences), followed by size exclusion chromatography (SEC) using a Superdex200 Increase 10/300 column (Cytiva), equilibrated in 10 mM Tris-HCl pH 7.8, 150 mM NaCl. SEC peak fractions containing spike protein were pooled, concentrated to 1 mg/ml, flash-frozen in liquid nitrogen and stored at −80°C.

### Cryo-EM sample preparation, imaging and processing

Wu spike (0.5 μM trimer) and W25 (1.7 μM) were incubated for 2 h on ice in a volume of 50 μl buffer containing 10 mM Tris-HCl pH 7.8 and 150 mM NaCl. 3.5 μl sample were adsorbed for 45 s on glow-discharged 400 mesh R1.2/R1.3 holey carbon grids (Quantifoil) and plunge-frozen in liquid ethane using a Vitrobot Mark IV (Thermo Fisher) with 1 s blotting time at 4°C and 99% humidity. Data were collected on a 300 kV Tecnai Polara cryo-EM (Thermo Fisher) equipped with a K2 Summit direct electron detector (Gatan) in super-resolution mode, at a nominal magnification of 31000×, with a pixel size of 0.63 Å/px and the following parameters: defocus range of −1 to - 2.5 μm, 50 frames per movie, 200 ms exposure per frame, electron dose of 6.2 e/Å^2^/s, leading to a total dose of 62 e/Å^2^ per movie. Data were collected using Leginon ^69^. Movies were aligned and dose-weighted using MotionCor2 ^70^. Initial estimation of the contrast transfer function (CTF) was performed with the CTFFind4 package via the Relion 3.0.7 GUI ^71,72^. Power spectra were manually inspected to remove ice contamination and astigmatic, weak, or poorly defined spectra. Subsequent image processing procedures are shown in Fig. S1. Particles were picked initially using the Relion 3.0.7 LoG autopicker. After 2D and 3D classification, the best defined 3D class was used as template for a second round of autopicking in Relion 3.0.7. 2D and 3D classification yielded 141.257 clean particles, which were further processed using Relion 3.0.7 or cryoSPARC 3.3.1^73^ as indicated in Fig. S1. Particles were subjected to an additional ab-initio reconstruction (3 classes, class similarity = 0), followed by a consensus refinement, exhibiting a 2 RBD-up, 1 RBD-down conformation. Subsequently, particles were locally refined to one RBD in the “up” conformation using a stalk/RBD/W25 mask, followed by a focused classification without alignment using a RBD/W25 mask for the identification of the most defined WuRBD/W25 complex subset. This particle subset was subjected to another local refinement using the RBD/NTD mask. Locally refined maps were used as reference in a heterogeneous refinement. This revealed that particles were partially in a 1 RBD-up, 2 RBD-down conformation. Clean particles were re-assigned to both global conformations, followed by non-uniform refinement, resulting in 3.8 Å map for the 2 RBD-up, 1 RBD-down class (map 1, Fig. S1) and a 3.81 Å map for the 1 RBD-up, 2 RBD-down class (map 2). Both particle subsets were pooled again and locally refined to the shared RBD in “up” conformation, resulting in a 5.92 Å map.

Omicron spike and W25 were mixed in 1:6 molar ratio and incubated on 37°C for 45 min. Omicron spike-W25 complex were adsorbed for 10 s onto glow discharged Quantifoil grids (Q1.2/1.3) and plunge frozen into liquid ethane using an EMGP2 system (Leica). Cryo-EM data collection was performed with Serial EM software (Version 3.1) using CRYO ARM 300 (JEOL) at the Center of Microscopy and microanalysis UQ. Movies were acquired in super resolution and CDS mode with a slit width of 10 eV using a K3 detector (Gatan) for 3.3 seconds during which 40 movie frames were collected with a 1 e-/Å^2^ per frame. Data were collected at a magnification of 60,000x, corresponding to a calibrated pixel size of 0.4 Å/pixel. Movies were binned 2x during motion Correction ^70^. A total of 9,258 micrographs were collected, 8,845 micrographs were selected based on an estimated resolution cutoff of 4.5 Å. Particles were extracted using Relion 3.0.7. All following processing steps were performed in Cryosparc 3.3.1 ^73^ as indicated in Fig. S3. Extracted particles were cleaned up from ice and false-positive picks using 2D classification. After 2D classification, most defined classes with high-resolution features were retained (164.806 particles) and used for an ab-initio 3D reconstruction (3 classes, class similarity = 0). Particles were further processed using heterogeneous refinement and previously generated maps, followed by a re-assignment to both global conformations from the Wuhan data set (maps 1 and 2, Fig. S1). A 3D variability analysis was performed for particles in 1 RBD-up, 2 RBD down conformation using a mask for the RBD in the “up” conformation. This revealed that particles were partially in a 3 RBD-down conformation. Using a less strict 2D class selection (222237 particles), the particle subset were then cleaned again using heterogeneous refinement, followed by re-assignment to the three identified global conformations. Particles subsets in the 2 RBD-up, 1 RBD-down and 1 RBD-up, 2 RBD-down conformation were cleaned in an additional round of heterogeneous refinement. The 1 RBD-up, 2 RBD-down particle subset showed the most defined RBD/W25 density and was therefore refined using non-uniform refinement (4.97 Å, map 4, Fig. S3), followed by local refinement using a RBD-up/NTD mask, resulting in a 6.04 Å map (map 5).

### Model building and refinement

The trimeric base, and the individual RBDs and NTDs of a Wu spike model (PDB 6ZXN,^63^) were fitted as rigid bodies in map 1 (Fig. S1). One RBD and the adjacent NTD, as well as an AlphaFold2 ^31,74^} model of the W25 nanobody were then fitted as rigid bodies in focus map 3 (Fig. S1), trimmed manually using Coot 0.9.5 ^75^, subjected to molecular dynamics flexible fitting using the Namdinator server ^76^ with standard parameters, followed by real space and ADP refinement in Phenix 1.19.2-4158 ^77^ and manual adjustment using Coot 0.9.5. For Omicron spike model building, a similar strategy was pursued, however using PDB 7WG6 ^78^ as starting model, and map 4 and 5 for fitting (Fig. S3). Molecular models and maps were visualized using the ChimeraX software ^79^.

### Surface plasmon resonance (SPR)

SPR measurements were performed using a Biacore 8K+ instrument (Cytiva). Purified W25-Fc, S309, DH1047 or C05 antibody were captured on a Protein A Series S Sensor Chip (Cytiva). Binding of SARS-CoV-2 spike variants were tested using a Multi-Cycle High Performance Kinetics assay. The antibodies were all prepared at 1 μg/mL in running buffer (HBS-P+, pH 7.4, Cytiva) and injected for 30 sec at 30 μL/min over Fc2 on all 8 channels. Multiple concentrations of SARS-CoV-2 spike variants were included: 0 nM, 1.40625 nM, 2.8125 nM, 5.625 nM, 11.25 nM, 22.5 nM, 45 nM and 90 nM. The proteins were injected over both Fc1 and Fc2 of Channels 1 −8 in series for 180 sec at 30 μL/min, followed by a dissociation period of 600 sec. The chip surface was regenerated between each cycle using 10 mM glycine pH 1.5. The data was double reference subtracted; reference cell subtraction (Fc1) from the active cell (Fc2), and zero concentration subtraction for each analyte-antibody pair. Sensorgrams for the association and dissociation phases were fitted to a 1:1 binding model, using Biacore Insight Evaluation software (Cytiva).

### ELISA

To test nanobody and antibodies, SARS-CoV-2 spike variant proteins in PBS pH7.4 were immobilised on Maxisorb ELISA (Nunc) plates at a concentration of 2 μg/mL overnight. Serial 5-fold dilutions of Fc-fusion nanobodies or antibodies in 1X KPL in blocking buffer (Seracare) were incubated with the immobilized antigen, followed by incubation with HRP-coupled anti-human IgG (MilleniumScience) and the chromogenic substrate TMB (ThermoFisher). Reactions were stopped with 2 M H_2_SO_4_ and absorption measured at 450 nm.

### Viruses

The SARS-CoV-2 Ancestral (Wu) variant, hCoV-19/Australia/QLD02/2020 (GISAID accession ID, EPI_ISL_407896), Alpha variant, hCoV-19/Australia/QLD1517/2021, (GISAID accession ID EPI_ISL_944644), Beta variant hCoV-19/Australia/QLD1520/2020, (GISAID accession ID EPI_ISL_968081), Delta: hCoV-19/Australia/QLD1893C/2021 (GISAID accession ID EPI_ISL_2433928) were kindly provided by Dr Alyssa Pyke (Queensland Health Forensic & Scientific Services, Queensland Department of Health, Brisbane, Australia). For Omicron BA.1 variant (hCoV-19/Australia/NSW-RPAH-1933/2021), Gamma variants, hCoV-19/Australia/NSW4318/2021, Lambda variants hCoV-19/Australia/NSW4431/2021 were kindly provided by A/Prof. Stuart Turville (University of New South Wales, Australia). Omicron BA.2 variant was kindly provided by Prof. Andreas Suhrbier (QIMR Berghofer Medical Research Institute). The passage 2 of SARS-CoV-2 variants (except Gamma and Lambda) was received, and passage 3 were propagated in TMPRSS2-expressing VeroE6 and used as virus stock. Passage 3 of and passage 4 of Gamma, Lambda and BA.2 variants were obtained and passage 4 was propagated in TMPRSS2-expressing VeroE6 and used as virus stock, viruses were stored at −80 °C. All working virus stocks were sequenced and validated using either Nanopore sequencing or Sanger sequencing on spike genes. For SARS-CoV-2 Beta variant (B.1.351) used in the animal experiment, was obtained from Andy Pekosz (Center for Emerging Viruses and Infectious Diseases (CEVID)). Infectious stocks were grown in Vero-Ace2-TMPRSS2 cells (BEI Resources) and stored at −80 °C. All work with infectious virus was performed in a biosafety level 3 laboratory and approved by the University of Queensland Biosafety committee (IBC/447B/SCMB/2021) and the Penn Institutional Biosafety Committee and Environmental Health and Safety.

### Plaque Reduction Neutralisation Test (PRNT)

The levels of neutralising antibodies were assessed using our established PRNT protocol ^28^ and supplementary materials and methods. Immuno-plaques were stained using anti-spike antibody (mouse CR3022) as the primary antibody for the following variants: SARS-CoV-2 Wu, Alpha, Gamma, Beta, Lambda, Kappa and Delta variants. Anti-M SARS-CoV-2 antibody was used as a primary antibody to stain the immune-plaques for Omicron variant. Immuno-plaques were analysed and counted using an automated foci counter program, Viridot ^80^.

### Cell-Cell fusion assay

Cell-cell fusion assay was conducted as previously described ^36^ and details are available in supplementary materials and methods. Briefly, HEK293T Lenti rLuc-GFP 1–7 (effector cells) were transfected with SARS-CoV-2 Spike glycoproteins of D614G, Beta and Omicron and human ACE2 plasmids were transfected into effector cells. The W25, W25-Fc and W25-FcM were incubated effector cells in 50 μl at 37 °C, 5% CO_2_ for 1 h, after which HEK293T Lenti rLuc-GFP 8–11 (target cells) were co-cultured to corresponding wells and incubated for 18–24 h, after which GFP-positive syncytia and *Renilla* luciferase were quantified. Negative controls (effector cells only, target cells only) and positive transfection controls (HEK293T Lenti rLuc-GFP 1–7 cells transfected with rLuc-GFP 8–11 plasmid) were always included. The assay was conducted separately for *Renilla* luciferase and GFP readout.

### Mouse infections

Animal studies were carried out in accordance with the recommendations in the Guide for the Care and Use of Laboratory Animals of the National Institutes of Health and the Guidelines to promote the wellbeing of animals used for Scientific purposes. The protocols were approved by the Institutional Animal Care and Use Committee at the University of Pennsylvania (protocol #807017) and the University of Queensland animal ethics committee (2021/AE000929 and 2021/AE001119). Heterozygous K18-hACE C57BL/6J mice (strain: 2B6.Cg-Tg(K18-ACE2)2Prlmn/J) were obtained from The Jackson Laboratory and/or bred in-house. Animals were housed in groups and fed standard chow diets. Virus inoculations were performed under anesthesia that was induced and maintained with ketamine hydrochloride and xylazine, and all efforts were made to minimize animal suffering. For prophylactic treatment study, mice of different ages and both sexes were infected with SARS-CoV-2 Beta variant at 1 × 10^3^ plaque forming units (PFU) via intranasal administration four hours after intraperitoneal administration of 100 ug W25-Fc or GFP-Nb-Fc in 100 ul PBS. For therapeutic treatment study, 5-6 weeks old female mice were infected with SARS-CoV-2 Beta (5 × 10^3^ FFU/mouse) or Omicron (1×10^5^ FFU/mouse) variants via intranasal administration. Infected mice were administrated with 100 ug W25-Fc or GFP-Nb-Fc via intraperitoneal (100 μl) or intranasal administration (20 μl) 24 hours post infection. Mice were monitored daily for weights and clinical respiratory disease scores were obtained. Mice were sacrificed when exhibiting greater than 20% weight loss or after reaching a respiratory disease score of 5 for longer than 2 days or greater than 5. Respiratory disease was scored based on activity, ruffled fur/piloerection, accelerated shallow breathing, labored breathing, tremor/tension/paralysis and/or weight loss of greater than 7.5% with a point added for each identified. Lungs and Nasal turbinates were collected on 2 or 4 days post-infection and viral titers were determined using plaque-forming assay, immunoplaque-forming assay and RT-PCR.

### Biodistribution of ^111^In-labelled W25-Fc in C57BL/6J mice

C57BL/6J female mice weighing 20-23 g (Janvier, France, La Rochelle) were housed in cages of 7-8 mice per cage. All mice were kept in a climate-controlled facility with a 12 hours light/dark cycle. The cages contained a biting stick, fed with commercial breeding diet ad libitum (1310 FORTI-Avlsfoder, Brogaarden, Altromin International) and had access to water. All procedures were conducted in accordance with the European Commission’s Directive 2010/63/EU, FELASA and ARRIVE guidelines for animal research and, with approval from The Danish Council for Animal Ethics (license numbers 2017-15-0201-01283) as well as the Department of Experimental Medicine, University of Copenhagen. Mice were evenly divided into six different groups. All mice were weighted prior to injection of ^111^In-W25Fc (1.2-1.5 MBq/mouse) through a tail vein. The animals were sacrificed under anesthesia (3% sevoflurane) by decapitation at the designated time points after tracer administration (group 1: 5 min, group 2: 20 min, group 3: 60 min, group 4: 3 hr, group 5: 5 hr and group 6: 24 hr). The mice were dissected and of each mouse the blood, lungs, liver, kidney, spleen, brain, muscle and tail were collected and placed into preweighed gamma counting tubes (Polypropylene, 5 mL 75×12 mm Ø, round base, Hounisens). The tubes with collected tissue were weighed and radioactivity measurements were conducted on a Cobra II Auto-Gamma Counter (Model D5005, Packard BioScience Company). The measurements were corrected for radioactivity decay and background. The ex vivo biodistribution results were expressed as a percentage of the injected radioactivity dose per gram of tissue (% ID/g) and nmol protein per tissue volume. For these calculations, tissue weight was converted to volume based on different tissue density for different organs. Radioactive counts in the tail were used as a quality control for the injections, which led to exclusion of one mouse in group 4 that had >20% ID in remaining in the tail.

## List of Supplementary Materials

Materials and methods

Fig. S1. Cryo-EM processing scheme for Wu spike/W25

Fig. S2. Conformational states of the Wu spike/W25 complex

Fig. S3. Cryo-EM processing scheme for Omicron spike/W25

Fig. S4. Detailed structural analysis of the Wu spike/W25 interaction, comparison to the binding modes of other nanobodies and of the ACE2 receptor

Fig. S5. Mutations in VOC spike proteins and characterisation of the spike preparations

Fig. S6. SPR sensorgrams of S309, DH1047 and C05

Fig. S7. ELISA binding curves of W25-Fc, EUA mAbs other epitope specific mAbs to SARS-CoV-2 Spikes variants

Fig. S8. Live virus neutralisation assay of W25 Nb against SARS-CoV-2 Wu and VOCs

Fig. S9. Amino acid residues on spike RBD involved in molecular contacts with W25

Fig. S10. Cell-cell fusion assay of SARS-CoV-2 spike

Fig.S11. Representative images illustrating GFP-positive syncytia from cell-cell fusion corresponding to Fig. S10

Fig. S12. Thermostability and nebulisation stability of W25

Fig.S13 Respiratory disease scores of K18-ACE2 mice infected with SARS-CoV-2 Beta variant and received W25-Fc 4 h prior to infection

Fig.S14. Pharmacokinetic of ^111^Indium radiolabelled W25-Fc

Fig.S15. Structural comparison of antibodies and nanobodies that induce unconventional neutralisation mechanism and mediate spike disruption

Table S1. Cryo-EM Data collection, processing and refinement statistics

Table S2. Survey of camelid and synthetic nanobody structures in complex with SARS-CoV2 Spike RBD

Movie S1. The movie illustrates the cryo-EM structure of the Wu spike/W25 complex, including a close-up of the RBD/W25 molecular model, highlighting the interaction interface.

## Acknowledgments

We thank Drs Matthias Flotenmeyer and Lou Brillault from Centre for Microscopy and Microanalysis (CMM), the University of Queensland for facilitating cryo-EM work to be carried out and for scientific and technical assistance. We thank UQBR animal staff (Maya Patrick and Barb Arnts) and UQBR facility at the Australian Institute for Bioengineering and Nanotechnology (AIBN). We thank Research Computing Centre (RCC), UQ for partially providing computational resources. We thank Queensland Health Forensic sand Scientific Services, Queensland and the Kirby Institute University of New Souths Wales for providing SARS-CoV-2 virus isolates. We thank the National Biologics Facility, AIBN for access to the Biacore 8K+ instrument for the SPR assay. We thank Prof. Andreas Suhrbier for providing Omicron BA.2 virus isolate. We thank Dr. Michael Landsberg for advice and helpful discussion. We thank Ian Mortimer (ITS Infrastructure Operations, UQ) for IT support. We thank the MPI-MG for granting access to the TEM instrumentation of the microscopy and cryo-EM service group. N.M would like to thank Alpaca ‘Budda’ whose nanobody was isolated from and Dzongsar Jamyang Khyentse Rinpoche for generous contribution.

## Author contributions

N.M.: Experimental design; conceptualised; designed, generated, purified spike proteins, monoclonal antibodies, W25-Fc and control Fc; characterised spike proteins and nanobodies; generated W25/Omicron spike dataset including sample preparation, optimisation and determination of optimal freezing conditions, image acquisition, performed Single Particle Analysis (SPA); performed PRNTs; performed K18-huACE2 mouse challenge experiments; generated figures; drafted and edited manuscript. S.M.L.: Performed Wu spike Hexapro/W25 and Omicron spike Hexapro/W25 cryo-EM single particle and structure analysis. A.A.A.: Propagated live virus stock, performed PRNTs, performed K18-huACE2 mouse challenge experiments and data analysis. P.W.: Performed K18-huACE2 mouse challenge experiment S.I.L.B.: Experimental design for the biodistribution studies, performed them with the help of J.T.J. and I.V.A. Y.S.L: Sample optimisation for W25/Omicron spike cryo-EM, determination optimal freezing conditions, data acquisition, data analysis. N.T.: Performed cell-cell fusion experiments. B.L.: Assisted in cloning and purification of spike proteins and antibodies; assisted in K18-huACE2 mouse challenge experiments. G.V.N.: Identification of W25 cloning of W25 and performed thermostability and nebulization stability assays of W25. J.J.: Performed SPA analysis and focuses classification for W25/Omicron dataset. D.P.: Cloned, expressed and purified published nanobodies. A.I.: Reformatted published nanobodies. J.D.S.: Generated live virus stock and generated VeroE6-TMPRSS6 cells for virus propagation. D.S.: Performed K18-huACE2 mouse challenge experiment. J.T.J.: Performed the biodistribution study with S.I.L.B and I.V.A. Y.C.: Performed optimization and purification of W25. J.B.: Assisted in Wu spike Hexapro/W25 cryoEM single particle and structure analysis. I.V.A.: Performed the biodistribution study with S.I.L.B and J.T.J. J.H.: Performed optimization and purification of W25 for biodistribution assays R.J.: Supervised vibrating mesh nebuliser stability assays. R.M.: Supervised vibrating mesh nebuliser stability assays. Z.M.C.: Performed optimization of W25 P.C.C supervise optimization of purification of W25 for cryo-EM and biodistribution. V.K.: Experimentally designed the biodistribution studies. T.M. and C.M.T.S.: Supervised Wu spike Hexapro/W25 cryo-EM single particle and structure analysis. A.A.K.: Obtained SARS-CoV-2 ancestral, Alpha, Beta isolates. Chief Investigator for BSL3 biocontainment facility at School of Chemistry and Molecular Biosciences. M.J.: Performed and analysed SPR experiment. A.K.: Designed the biodistribution studies, supervised the work of J.T.J. M.M.H.: Designed the labelling and biodistribution studies, supervised the work of I.V.A. and S.I.L.B. K.A.J.: Supervised K18-huACE2 prophylactic mouse work. D. Sc.: Prepared and purified Wu Hexapro spike, prepared Wu spike Hexapro/W25 complex cryo-EM sample, performed and supervised Wu spike Hexapro/W25 cryo-EM and Omicron spike Hexapro/W25 single particle and structure analysis, generated figures, wrote initial manuscript draft. D.W.: Performed SPA of W25/Omicron dataset. The overall project was supervised, conceptualized and edited by D.W. A.R.F. and D.Sc.

## Data and materials availability

All data are available in the main text or the supplementary materials. Cryo-EM maps and molecular models have been deposited in the Electron Microscopy Data Bank (EMDB) and the Protein Data Bank (PDB), respectively, with accession codes EMD-XXXX (map 1), EMD-XXXX (map 2), EMD-XXXX (map 3), EMD-XXXX (map 4), EMD-XXXX (map 5) and PDB XXXX (Wu NTD/RBD/W25) and XXXX (Omicron NTD/RBD/W25).

