## Supplementary for "An alpaca-derived nanobody recognizes a unique conserved epitope and retains potent activity against the SARS-CoV-2 omicron variant"

### Materials and Methods

*Expression and purification of SARS-CoV-2 Spike*

Soluble, trimeric spikes (residue 1-1204 amino acid) of SARS-CoV 2 /human/China/Wuhan-Hu-1/2019 (referred as Wu) (Genbank: MN908947), SARS-CoV-2 Beta variant (B.1.351) (L18F, D80A, D215G, Δ242-244, K417N, E484K, N501Y, D614G and A701V), Alpha variant (B.1.1.7) (Δ69-70, ΔY144-145, N501Y, A570D, D614G, P681H, T716I, S982A and D1118H), Gamma variant (P.1) (L18F, T20N, P26S, D138Y, R190S, K417T, E484K, N501Y, D614G, D655Y, T1027I and V1176F) , Lambda variant (C.37) (G75V, T76I, R246N, Δ247-253, L452Q, F490S, D614G and T859N), Delta variant (B.1.617) T19R, E156G, Δ157-158, L452R, T478K, D614G, P681R and D950N, Omicron (BA.1) (A67V, Δ69-70, T95I, G142D, Δ143-145, Δ211, L212I, 214EPEinsert, G339D, S371L, S373P, S375F, K417N, N440K, G446S, S477N, T478K, E484A, Q493R, G496S, Q498R, N501Y, Y505H, T547K, D614G, H655Y, N679K, P681H, N796Y, N856K, Q954H, N969K, L981F) and Omicron (BA.2) (T19I, L24S, Δ25-26, A67V, Δ69-70, G142D, V213G, S371F, S373P, S375F, T376A, D405N, R408S, K417N, N440K, S477N, T478K, E484A, Q493R, Q498R, N501Y, D614G, H655Y, N679K, P681H, N764K, D796Y, Q954H and N969K) spike mutations were added *in silico* into the codon-optimised Wuhan reference stain and were cloned to pNBF plasmid. Spike proteins contain 6 proline mutations (F817P, A892P, A899P, A942P, K986P and V987P) and substituted at the furin cleavage site (residues 682–685).

*Nanobodies expression and purification*

The pHen6-W25 vector inoculated in 20 mL of liquid LB-medium containing 100 µg/ml ampicillin and 1% glucose. The bacteria were cultured at 37 °C with agitation for 16 h. The bacteria were then diluted into 1L Terrific Broth (TB) medium containing 100 µg/ml ampicillin, 2 mM MgCl2, 0,1% glucose and incubated at 37 °C to an OD600 of 0.6–0.9. The expression of the nanobodies was induced by adding 1 mM of IPTG (isopropyl-β-d-1-thiogalactopyranoside) for 20 h at 28 °C. Bacteria were collected by centrifugation at 8000 rpm for 8 min at 4 °C. The harvested bacteria were resuspended in a 12 mL TES buffer (0.2 M Tris pH 8.0, 0.5 mM EDTA, 0.5 M sucrose) and incubated for 1 h on ice, then incubated for another hour on ice in 18 mL TES buffer, diluted 4 times and centrifuged at 8000 rpm at 4 °C to pellet down cell debris. The supernatant was loaded on 5 mL of HisPur Ni–NTA agarose resin which was pre-equilibrated with binding buffer (Tris 50 mM, NaCl 500 mM, imidazole 10 mM pH 7.5). The lysed cells containing His- and myc-tagged nanobodies were added to the column followed by adding the column's volume in binding buffer for a total of eight times. The column was washed by adding eightfold the column´s volume with wash buffer (Tris 50 mM, NaCl 500 mM, imidazole 30 mM pH a 7.5), and eluted with 15 mL of elution buffer (Tris 50 mM, NaCl 150 mM, 150 mM imidazole, 1 mM DTT pH 7.5).

**Plaque Reduction Neutralisation Test (PRNT)**

The levels of neutralising antibodies were assessed using our established PRNT protocol [1]. Briefly, purified mAbs were five-fold serially diluted in DMEM containing 2% HI-FCS and 1% P/S (Gibco). Subsequently, serial diluted antibodies were incubated with 50-100 immuno-plaques of SARS-CoV-2 and incubated at 37 °C for 1 h. Then, 50 µL of mixture (virus/antibody complex) was added onto pre-seeded VeroE6 cells in 96-well plates at 6 x$10^4$ cells/well and incubated at 37 °C for 30 min. Following, overlay medium (containing 1% CMC, 1X M199, 2% HI-FCS and 1% (P/S)) was added on top of the inoculum and incubated at 37 °C with 5% of $CO_2$. Twenty-four hours after infection, the overlay was removed, and the monolayer was fixed with cold fixative solution (80% acetone and 20% phosphate buffered saline (PBS)) and incubated at -20°C for 30min. Fixative reagent was removed, and the monolayer was fully dried. Monolayer was then treated with 100 µl of 1X milk blocking solution (KPL, Seracare) diluted in 1X-PBS containing 0.05% Tween 20 (PBS-T) and incubated at 4°C for overnight. Next, the blocking buffer was removed, and immuno-plaques were stained using anti-spike antibody (mouse CR3022) as the primary antibody for the following variants: SARS-CoV-2 Wu, Alpha, Gamma, Beta, Lambda, Kappa and Delta variants. Anti-M SARS-CoV-2 antibody was used as a primary antibody to stain the immune-plaques for Omicron variant. The monolayer was incubated at 37 °C for 1 h with the primary antibody and the unbounded antibody was removed by five consecutive washes with 5min incubation between each washed. To reveals the immuno-plaques, an infra-red dye conjugated secondary antibody (IRDye 800CW Goat anti-Mouse or Goat anti-streptavidin, MillenniumScience) was added and incubated at 37 °C for 1 h, followed by five consecutive washes with 5min incubation in between. Finally, plate was fully dried avoiding long light exposure and scanned using the LI-COR Biosciences Odyssey Infrared Imaging System (Odessey CLx, Li-COR, USA). Immuno-plaques were analysed and counted using an automated foci counter program, Viridot [2]. A stock of primary and secondary antibodies at 1 mg/mL were diluted 1/1000 and 1/2500 in blocking buffer solution, respectively and 50 µl/well was used for the staining.

**Cell-cell fusion assay**

HEK293T Lenti rLuc-GFP 1–7 (effector cells) and HEK293T Lenti rLuc-GFP 8–11 (target cells) were seeded separately at $7.5\times10^5$ per well in a six-well dish in 3 ml of PRF-DMEM-10% and incubated overnight at 37 °C, 5 % $CO_2$. Transfection mixes were set up in 200 µl Opti-MEM (Gibco) with the *Trans*IT-X2 Dynamic Delivery System as per the manufacturer's recommendations (Mirus). SARS-CoV-2 Spike glycoproteins of D614G, Beta and Omicron and human ACE2 plasmids were transfected into effector cells. A mock-transfected (pcDNA3.1 empty plasmid, - vGP) and positive transfection control (250 ng rLuc-GFP 8–11 plasmid) was also set up. W25, its derivatives and control antibody were diluted to specific concentration in sterile 1.5 ml tubes using serum-free PRF-DMEM and plated at 25 µl/well in a white-bottomed, sterile 96-well plate (Corning), including no antibody controls. The antibodies were incubated with $2\times10^4$ effector cells in 50 µl at 37 °C, 5% $CO_2$ for 1 h, after which target cells were co-cultured to corresponding wells and incubated for 18–24 h, after which GFP-positive syncytia and *Renilla* luciferase were quantified. Negative controls (effector cells only, target cells only) and positive transfection controls (HEK293T Lenti rLuc-GFP 1–7 cells transfected with rLuc-GFP 8–11 plasmid) were always included. The assay was conducted separately for *Renilla* luciferase and GFP readout.

To quantify *Renilla* luciferase expression in fusion assays media were replaced with 100 µl of phosphate-buffered saline (PBS) followed by 60 µl of diluted substrate, Coelenterazine-H, 1 µM (Promega) 1:400 with PBS. The plate was incubated in the dark for 2 min then read on the GloMax Multi$^+$ Detection System (Promega).

To quantify GFP expression, cells were plated in clear flat-bottomed 96-well plates (Nunc) and imaged every hour using the IncuCyte S3 live cell imaging system (Essen BioScience). Five fields of view were taken per well at 10× magnification, and GFP expression was determined using the total integrated intensity metric included in the IncuCyte S3 software (Essen BioScience). To analyse images generated on the IncyCyte S3, a collection of representative images is first taken to set fluorescence and cellular thresholds, which allows for the removal of background fluorescence, and selection of cell boundaries ('objects') by creating 'masks'. Following this, the total integrated intensity metric can be accurately calculated by the software, which takes the total sum of objects' fluorescent intensity in the image, expressed as green count units (GCU) µm.

**Plaque-forming assay and immunoplaque-forming assay**

Tissues were weighed and homogenized with zirconia beads in a FastPrep-24 instrument (MP Biomedicals) in 0.6 ml of DMEM media supplemented with 10% heat-inactivated FBS. Tissue homogenates were clarified by centrifugation at 10,000 r.p.m. for 5 min and stored at −80 °C. Vero-Ace2-Tmprss2 cells (BEI Resources) were seeded at a density of $2.5 \times 10^6$ cells per plate in flat-bottom 6-well tissue culture plates. The following day, media was removed and replaced with 200 µl of tenfold serial dilutions of the tissue homogenate to be titered, diluted in DMEM. One hour later, seaplaque/seakem agarose was added. Plates were incubated for 72 h, then fixed with 4% paraformaldehyde in phosphate-buffered saline for at least 1 hour. Plaques were visualized with 0.05% (wt/vol) crystal violet in 20% methanol and washed with water prior to enumeration of plaques.

**Viral load via RT-qPCR or immuno-plaque assay (iPA)**

For RT-qPCR, tissue homogenate was added to Trizol at 1:3 ratio and extracted using Zymo Direct-zol RNA kit following manufactures protocol. RNA was reverse transcribed and amplified using iScript cDNA Synthesis Kit (Biorad). Gene-specific primers for SARS-CoV-2 N gene (F:TTACAAACATTGGCCGCAAA & R:GCGCGACATTCCGAAGAA) and mouse HPRT1 (F: GTTGGATACAGGCCAGACTTTGTTG & R: GAGGGTAGGCTGGCCTATTGGCT) with Power SYBR Green PCR Master Mix (Applied Biosystems) were used to amplify viral and cellular RNA by QuantStudio 3 Flex Real-Time PCR Systems (Applied Biosystems). The relative expression levels of target genes were calculated using the standard curve method using quantitative synthetic RNA (ATCC) and normalized to HPRT1 RNA as an internal control.

For iPA, organs such as lung and nasal turbinates were collected in preweighted tubes and homogenised using a tissue homogenizer (TissueLyser LT, Qiagen). Homogenates were then clarified by centrifugation at 10000 × g, 4 °C for 5 minutes. Supernatants were collected and stored at −80°C. Viral loads were determined by iPA on VeroE6 following our optimised protocol for viral titration. Viral titres were expressed in FFU per grams of tissue (FFU/g).

**Radiochemistry**

[$^{111}$In]$InCl_3$ was purchased from Curium. The analytical-HPLC system consists of a 170U UVD detector, a Scansys radiodetector and a Dionex system connected to a P680A pump. The system was run by Chromeleon software. The radiochemical conversion (RCC) of the radiolabeled compounds was determined by analyzing an aliquot of the crude reaction mixture by radio-HPLC analysis integrating the radioactive peaks of the chromatogram [3]. Radiolabeled products were characterized by associating the UV-HPLC traces of the authentic cold compounds with the radio-HPLC chromatogram of the reaction mixtures. The radiochemical yield (RCY) was determined using the initial activity at the beginning of the synthesis and that of the formulated product at the end of thes, corrected for decomposition and corrected for decay.

**$^{111}$In-labeling DOTA-tetrazine**

1,4,7,10-tetraazacyclododecane-1,4,7,10-tetraacetic acid (DOTA) -PEG$_{11}$-tetrazine was dissolved (2 mg/mL) in metal-free water and stored at -80°C before use. In general, an aliquot of 50-100 µL (10-30 MBq) of [111In]indium chloride in 0.1M HCl was combined with 2 µL DOTA-PEG11-tetrazine and 1M $NH_4OAc$ buffer (pH 5.5) at a volume ratio of 1:10 was added. The mixture was shaken at 600 rpm for 5 min at 60 °C in an Eppendorf ThermoMixer C. Then, 10 mM diethylenetriamine-pentaacetic acid (DTPA, volume ratio 1:10) and 2 µL 10 mg/mL gentisic acid in saline was added and the solution was shaken for an additional 5 min at 60 °C in an Eppendorf ThermoMixer C. Quantitative labeling yield and a radiochemical purity reater than 95% were obtained with this method, as confirmed by radio-HPLC.

**Trans–cyclooctene (TCO)-modifications W25-Fc**

W25-Fc (2.0 mg/mL) in PBS (pH 7.4) was aliquoted in five vials of 100 µL. To each aliquot, 25 eq. TCO-PEG4-NHS (Broadpharm, BP-22418) and sodium carbonate buffer (1 M, 3.1 µL, pH 8.0) was added. The mixture was shaken at 600 rpm for 2 hours at room temperature in the dark. Unreacted TCO-PEG4-NHS was removed by purification with Zeba spin desalting columns (7K MWCO, 0.5 mL, 89882, Thermofisher) and eluted in PBS (pH 7.4), with >95% protein recovery after purification. Final protein concentration was 1.7 mg/mL, measured with NanoDrop (NanoDrop 2000, ThermoScientific). Titration experiments were conducted to quantify the amount of reactive TCOs per protein-conjugate. $^{111}$In-labeled Tz stock was diluted accordingly to add 5 µL an excess of $^{111}$In-labeled Tz (2 eq. of Tz per protein-conjugate) to 5 µL of TCO-HSA and the mixture was shaken at 600 rpm for 1 hour at 37 °C. 3 µL NuPAGE™ LDS Sample Buffer (NP0007, Invitrogen) was added and the mixture was shaken for 10 minutes at 70 °C. Samples were applied to NuPAGE™ 4 to 12%, Bis-Tris, 1.0 mm, Mini Protein Gel, 12-well (NP0322BOX, Invitrogen) SDS-PAGE gels. SDS-PAGE gels were exposed to phosphor storage screens and read by a Cyclone Storage Phosphor System (PerkinElmer Inc.). Quantification of plate readings was done with Optiquant software (version

5.00, PerkinElmer Inc.). Quantification by radioactive SDS-PAGE revealed presence of approximately 1.6 reactive TCO/protein.

**Radiolabelling of TCO-W25-Fc with $^{111}$In-tetrazine**

To a 5 mL Eppendorf vial was added TCO-W25-Fc (465 µL, 10.16 nmol protein, 16.26 nmol TCO) and $^{111}$In-Tz (68 MBq, 95 µL, 1.6 nmol, 0.1 TCO/Tz eq.). The mixture was shaken at 600 rpm for 1 hour at 37 °C, giving a radiochemical conversion of 68%. Ultrafiltration (Vivaspin 500, 5,000 MWCO PES, Sartorius) yielded in 40 MBq $^{111}$In-W25-Fc with a radiochemical purity of 96%, determined by radio-HPLC. Protein concentration after purification was measured with NanoDrop (NanoDrop One, ThermoScientific). The $^{111}$In-labeled protein was formulated in PBS to a final concentration of 15 MBq/mL.

**Table S1. Cryo-EM Data collection, processing and refinement statistics**

**Wu spike/W25 complex**

| | |
|---|---|
| Magnification | 31000 |
| Voltage | 300 |
| Total dose per movie stack [e/Å$^2$] | 62 |
| Defocus range [$\mu$m] | 0.5-2.5 |
| Pixel size [Å] | 0.625 |
| Initial particle no. | 284399 |
| Final particle no. | 76384 (map 1), 56250 (map 2), 132634 (map 3) |
| Map resolution [Å] | 3.80 (map 1), 3.81 (map 2), 5.92 (map 3) |
| FSC threshold | 0.143 |
| Map sharpening B factor [Å$^2$] | -96.4 (map 1), -94.5 (map 2), -484.1 (map 3) |

| | Map 3 |
|---|---|
| Starting model | PDB 6dsz |
| Model fit CCs - mask/box/peaks/volume | 0.70/0.82/0.83/0.71 |
| Model resolution - masked/unmasked | 6.7/6.6 (5.0/5.1) |
| FSC threshold | 0.5 (0.143) |
| Model composition | |
| Protein residues | 542 |
| Ligands | BMA:3, NAG:9 |
| Average B factors [Å$^2$] | |
| Protein | 269.6 |
| Ligand | 329.6 |
| R.m.s. deviations | |
| Bond lengths [Å] | 0.003 |
| Bond angles [°] | 0.72 |
| Validation | |
| MolProbity score | 2.12 |
| Clashscore | 6.27 |
| Poor rotamers [%] | 1.5 |
| Ramachandran plot | |
| Favoured [%] | 85.9 |
| Allowed [%] | 14.1 |
| Disallowed [%] | 0 |

**Omicron spike/W25 complex**

| | |
|---|---|
| Magnification | 60000 |
| Voltage | 300 |
| Total dose per movie stack [e/Å$^2$] | 40 |
| Defocus range [$\mu$m] | 0.5-2.5 |
| Pixel size [Å] | 0.4 |
| Initial particle no. | 222237 |
| Final particle no. | 32512 (map 4 and 5) |
| Map resolution [Å] | 4.97 (map 4), 6.04 (map 5) |
| FSC threshold | 0.143 |
| Map sharpening B factor [Å$^2$] | -189.6 (map 4), -411.8 (map 5) |

| | Map 5 |
|---|---|
| Starting model | PDB 7wg6 |
| Model fit CCs - mask/box/peaks/volume | 0.77/0.91/0.91/0.77 |
| Model resolution - masked/unmasked | 6.2/6.2 (4.9/4.9) |
| FSC threshold | 0.5 (0.143) |
| Model composition | |
| Protein residues | 567 |
| Ligands | BMA:2, NAG:9, MAN:2 |
| Average B factors [Å$^2$] | |
| Protein | 325.7 |
| Ligand | 318.3 |
| R.m.s. deviations | |
| Bond lengths [Å] | 0.003 |
| Bond angles [°] | 0.659 |
| Validation | |
| MolProbity score | 2.22 |
| Clashscore | 18.16 |
| Poor rotamers [%] | 0 |
| Ramachandran plot | |
| Favoured [%] | 92.7 |
| Allowed [%] | 7.1 |
| Disallowed [%] | 0.2 |

**Table S2. Survey of camelid and synthetic nanobody structures in complex with SARS-CoV2 Spike RBD**

| NB | Type | Method | PDB ID | Reference PMID |
|---|---|---|---|---|
| C1 | side-1 | X-ray | 7oap | 34552091 |
| C5 | top | X-ray | 7oao | 34552091 |
| H3 | top | X-ray | 7oap | 34552091 |
| F2 | side-1 | X-ray | 7oay | 34552091 |
| Nb17 | side-2 | Cryo-EM | 7mej | 34344900 |
| Nb21 | top | Cryo-EM | 7mdw | 34344900 |
| Nb36 | side-2 | Cryo-EM | 7mej | 34344900 |
| Nb105 | side-1 | Cryo-EM | 7mdw | 34344900 |
| VHH E | top | Cryo-EM | 7ksg, 7b17 | 33436526 |
| VHH V | side-1 | Cryo-EM | 7b11, 7b17 | 33436526 |
| VHH U | side-1 | X-ray | 7kn5 | 33436526 |
| VHH W | side-1 | X-ray | 7kn7 | 33436526 |
| Ty-1 | top | Cryo-EM | 6zxn | 32887876 |
| H11-D4 | top | X-ray | 6yz5 | 32661423 |
| H11-H4 | top | X-ray | 6zbp | 32661423 |
| NM1226 | side-1 | X-ray | 7nkt | 33904225 |
| NM1230 | top | X-ray | 7b27 | 33904225 |
| WNb-2 | top | X-ray | 7ldj | 33893175 |
| WNb-10 | side-1 | Cryo-EM | 7lx5 | 33893175 |
| Re5D06 | top | X-ray | 7olz | 34302370 |
| Re9F06 | side-1 | X-ray | 7olz | 34302370 |
| Nb20 | top | X-ray | 7jvb | 33154108 |
| Nanosota-1 | top | X-ray | 7km5 | 34338634 |
| MR17 (synthetic) | top | X-ray | 7c8w | 34330908 |
| SR4 (synthetic) | top | X-ray | 7c8v | 34330908 |
| SR31 (synthetic) | side-1 (unconventional) | X-ray | 7d2z | 33657135 |
| Sb16 (synthetic) | top | X-ray | 7kgk | 34537245 |
| Sb45 (synthetic) | top | X-ray | 7kgj | 34537245 |
| Sb 68 (synthetic) | side-1 | X-ray | 7klw | 34537245 |

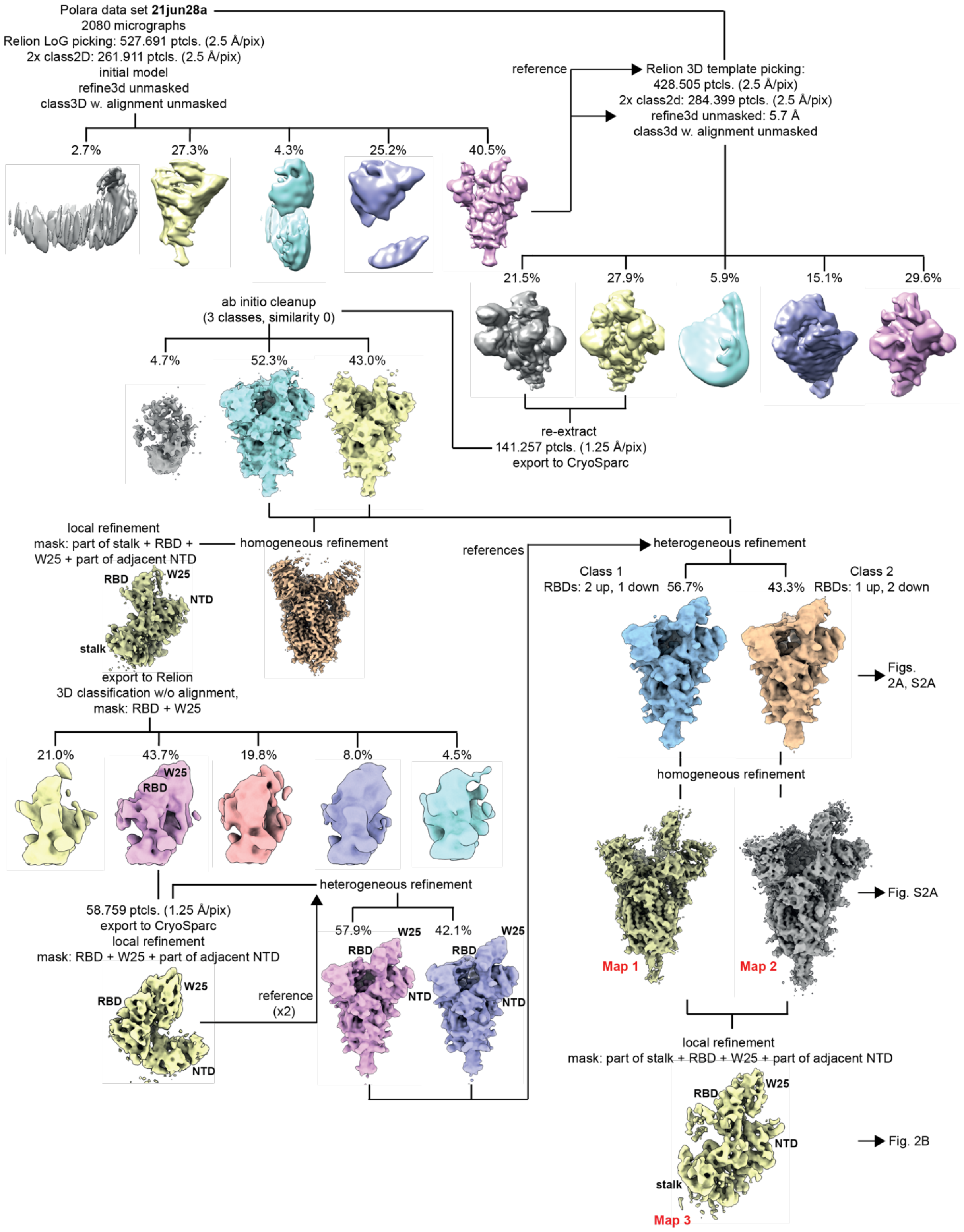

Polara data set **21jun28a**
2080 micrographs
Relion LoG picking: 527.691 ptcls. (2.5 Å/pix)
2x class2D: 261.911 ptcls. (2.5 Å/pix)
initial model
refine3d unmasked
class3D w. alignment unmasked
2.7%
27.3%
4.3%
25.2%
40.5%
reference
Relion 3D template picking:
428.505 ptcls. (2.5 Å/pix)
2x class2d: 284.399 ptcls. (2.5 Å/pix)
refine3d unmasked: 5.7 Å
class3d w. alignment unmasked
21.5%
27.9%
5.9%
15.1%
29.6%
ab initio cleanup
(3 classes, similarity 0)
4.7%
52.3%
43.0%
re-extract
141.257 ptcls. (1.25 Å/pix)
export to CryoSparc
local refinement
mask: part of stalk + RBD +
W25 + part of adjacent NTD
homogeneous refinement
references
heterogeneous refinement
RBD
W25
NTD
stalk
Class 1
RBDs: 2 up, 1 down 56.7%
Class 2
43.3% RBDs: 1 up, 2 down
Figs.
2A, S2A
export to Relion
3D classification w/o alignment,
mask: RBD + W25
21.0%
43.7%
19.8%
8.0%
4.5%
W25
RBD
homogeneous refinement
Map 1
Map 2
Fig. S2A
58.759 ptcls. (1.25 Å/pix)
export to CryoSparc
local refinement
mask: RBD + W25 + part of adjacent NTD
heterogeneous refinement
57.9%
W25
42.1%
W25
RBD
RBD
NTD
NTD
reference
(x2)
W25
RBD
NTD
local refinement
mask: part of stalk + RBD + W25 + part of adjacent NTD
W25
RBD
NTD
stalk
Map 3
Fig. 2B

Map 1

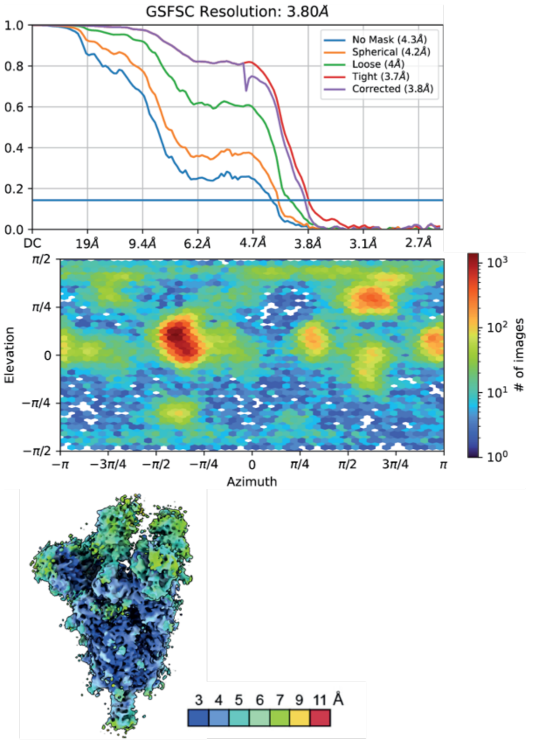


Map 2

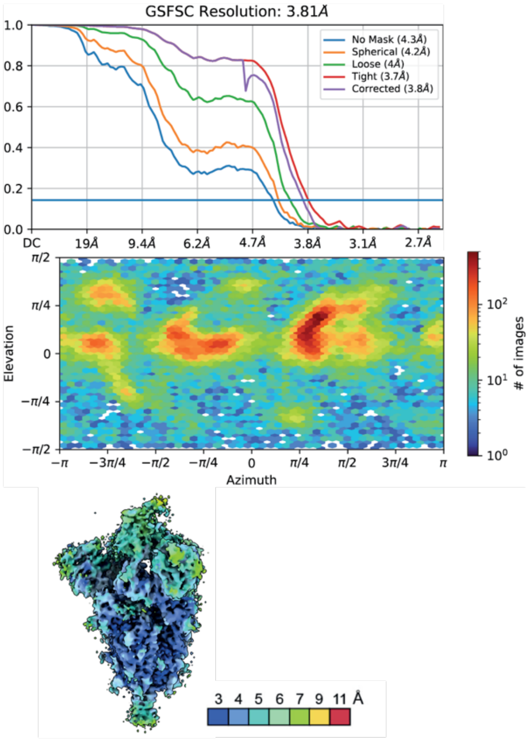


Map 3

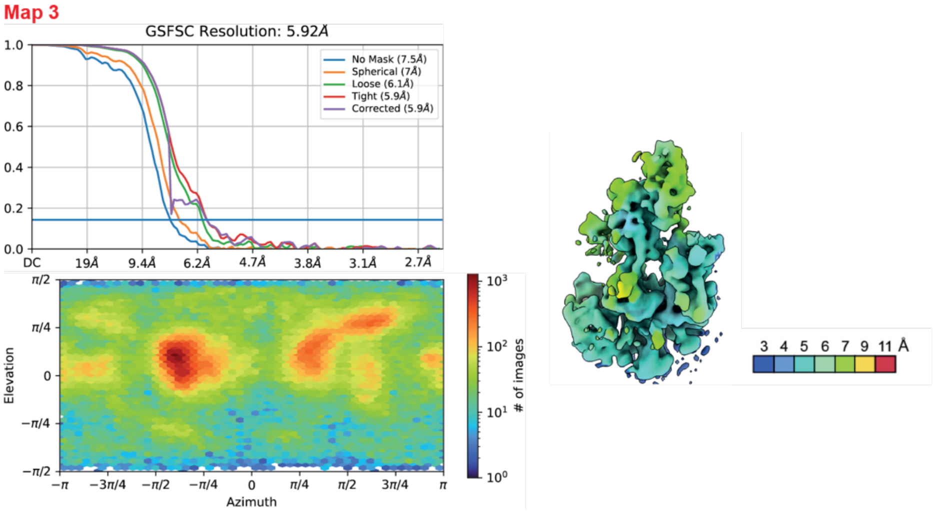


**Fig. S1. Cryo-EM processing scheme for Wu spike/W25**

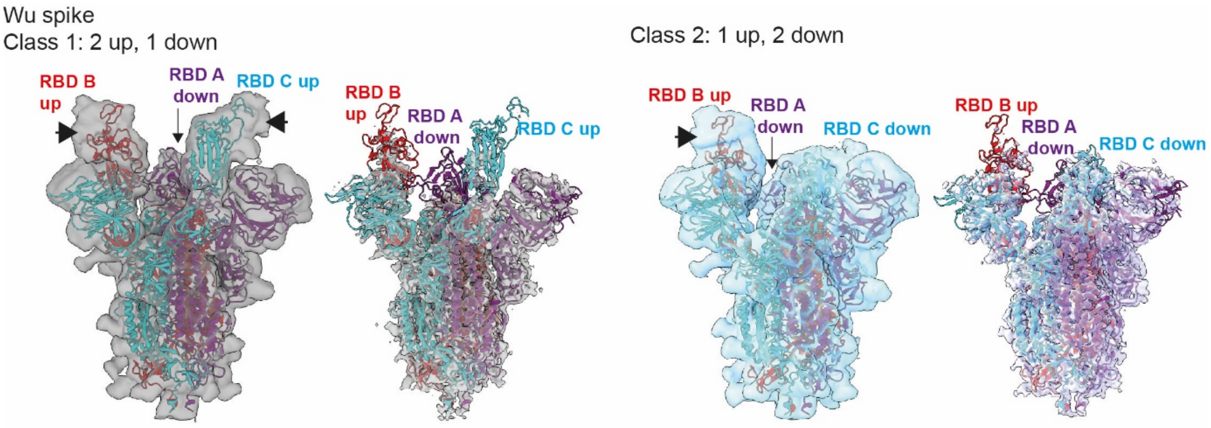


**Fig. S2. Conformational states of the Wu spike/W25 complex.**
Cryo-EM densities representing spike conformational class 1 (grey – map 1 [Fig. S1]) and class 2 (cyan, map 2 [Fig. S1]. The left panel for each class shows a filtered map at higher contour level to better illustrate the extra densities corresponding to W25 (indicated by the black arrows). The spike protein is represented as in Fig. 2A

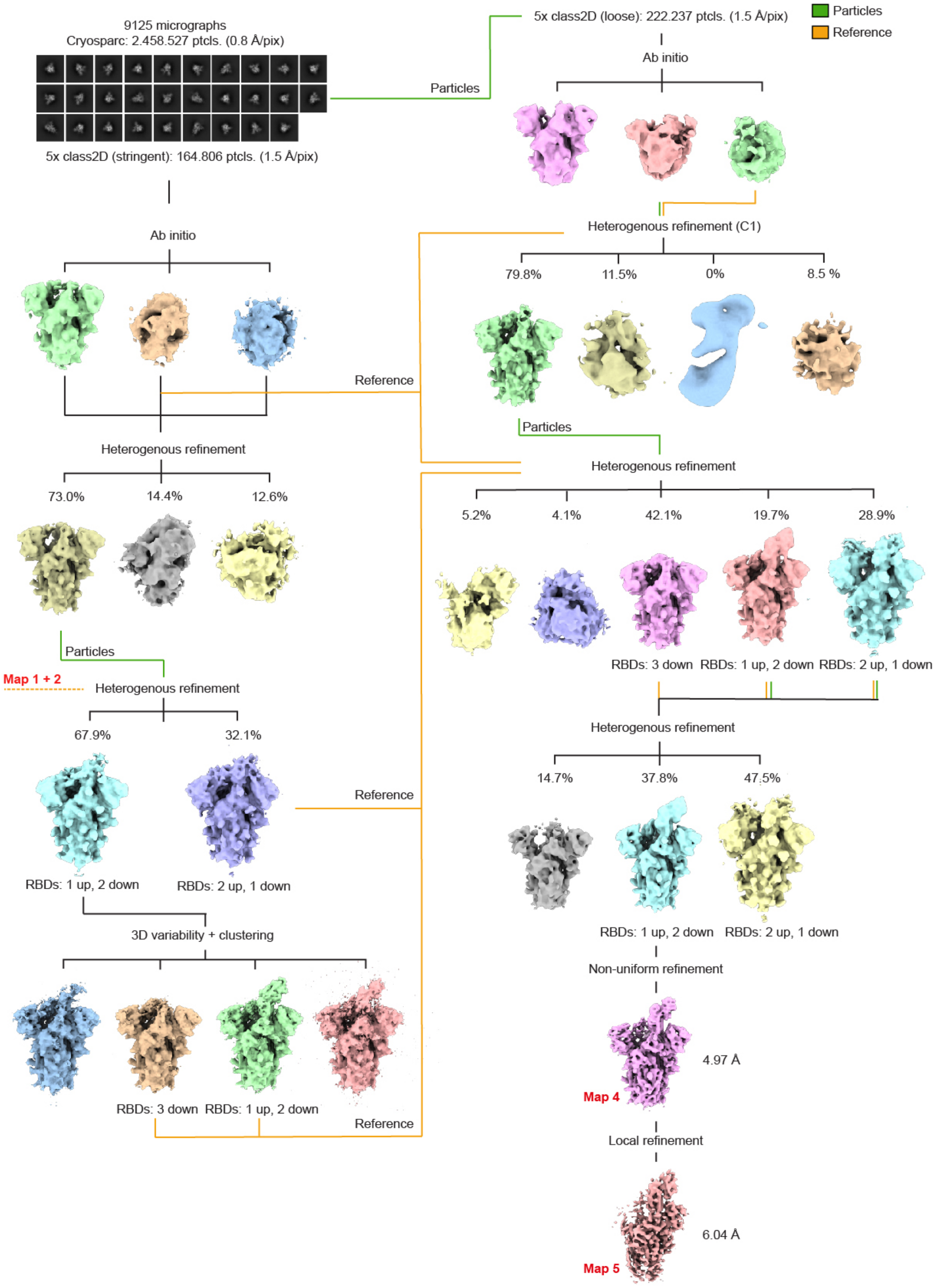

9125 micrographs
Cryosparc: 2.458.527 ptcls. (0.8 Å/pix)
5x class2D (stringent): 164.806 ptcls. (1.5 Å/pix)
Particles
5x class2D (loose): 222.237 ptcls. (1.5 Å/pix)
Particles
Reference
Ab initio
Ab initio
Heterogenous refinement (C1)
79.8%
11.5%
0%
8.5 %
Reference
Heterogenous refinement
73.0%
14.4%
12.6%
Particles
Heterogenous refinement
5.2%
4.1%
42.1%
19.7%
28.9%
RBDs: 3 down
RBDs: 1 up, 2 down
RBDs: 2 up, 1 down
Particles
Map 1 + 2
Heterogenous refinement
67.9%
32.1%
Reference
RBDs: 1 up, 2 down
RBDs: 2 up, 1 down
Heterogenous refinement
14.7%
37.8%
47.5%
RBDs: 1 up, 2 down
RBDs: 2 up, 1 down
3D variability + clustering
Non-uniform refinement
4.97 Å
Map 4
RBDs: 3 down
RBDs: 1 up, 2 down
Reference
Local refinement
6.04 Å
Map 5

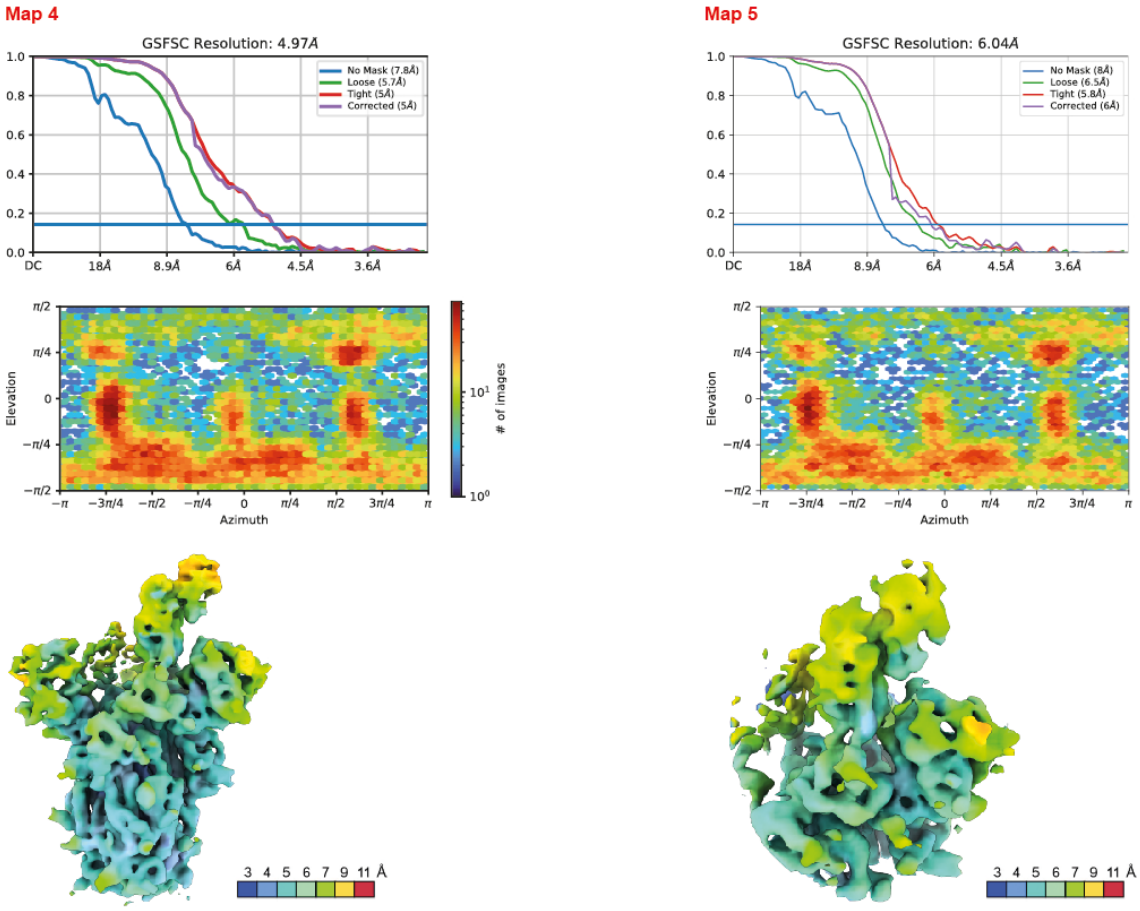


**Fig. S3. Cryo-EM processing scheme for Omicron spike/W25**

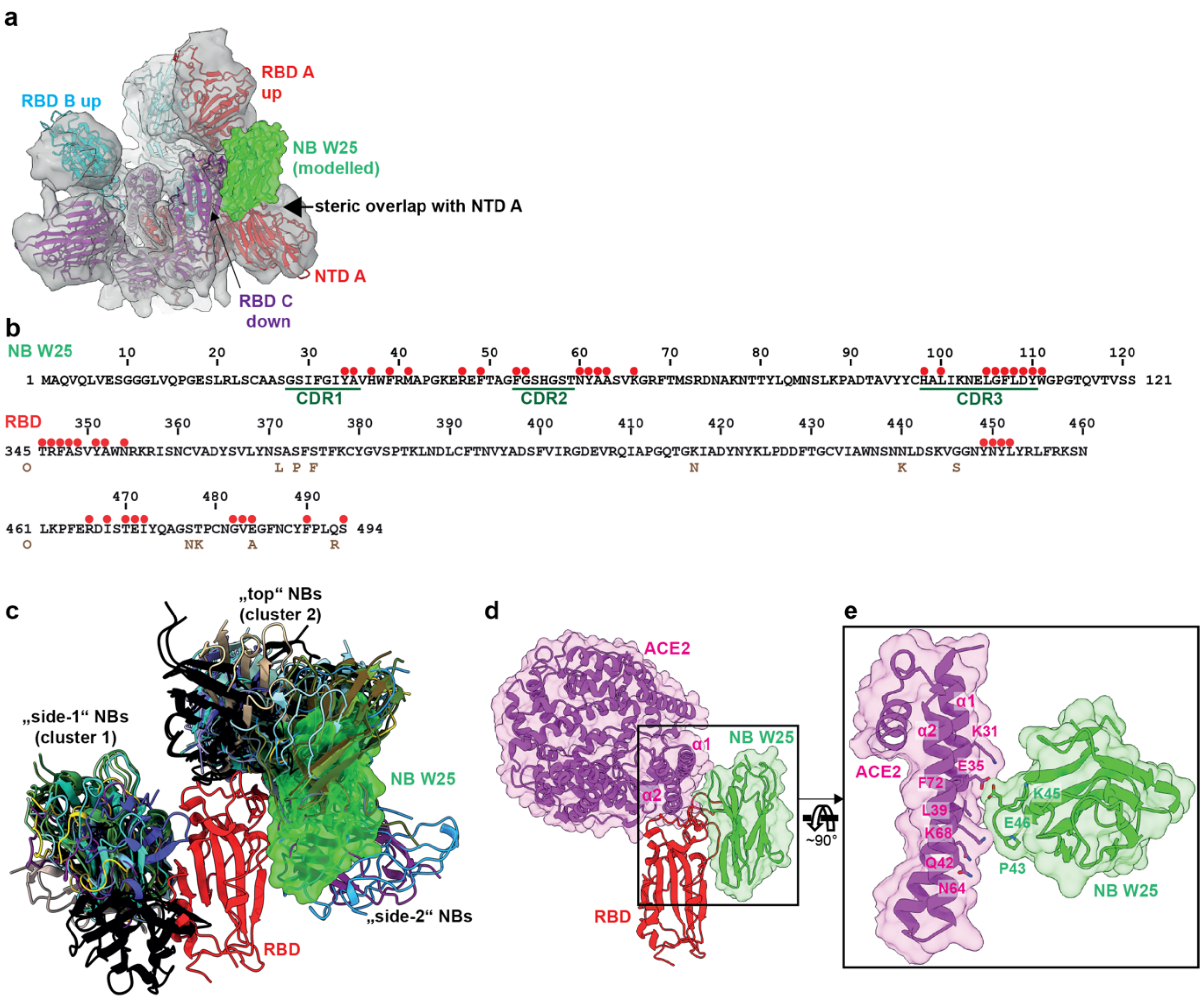


**Fig. S4. Detailed structural analysis of the Wu spike/W25 interaction, comparison to the binding modes of other nanobodies and of the ACE2 receptor.**

**a** Superposition of a W25-bound RBD with the “down”-RBD of the spike trimer. A steric clash between the modelled W25 position and the adjacent NTD is indicated by the arrow.

**b** Upper panel: amino acid sequence of W25. All amino acid residues involved in molecular contacts with the spike RBD, including main chain interactions, are indicated as red dots above the sequence. CDRs are underlined. Lower panel: Amino acid sequence of spike RBD (original Wuhan isolate). All amino acid residues involved in molecular contacts with the spike RBD are indicated as red dots above the sequence.

**c** Superposition of all camelid and synthetic nanobody/SARS-CoV2 RBD structures available in the PDB (Table S1). Only the RBD of PDB entry 7oap (red cartoon) is shown for clarity. Nanobodies are shown in cartoon representation, and nanobody W25 additionally as semi-transparent surface.

**d** Superposition of the spike RBD/ACE2 receptor complex (PDB 7a94,[4] with the RBD/nanobody W25 structure. For clarity, only the RBD of the W25 complex is shown as red cartoon. ACE2 (magenta) and W25 (green) are depicted as cartoons together with a semi-transparent representation of their solvent-accessible surfaces.

**e** Detailed view of the structural superposition from H. Selected ACE2 (magenta) and W25 (green) amino acid side chains are shown in stick representation.

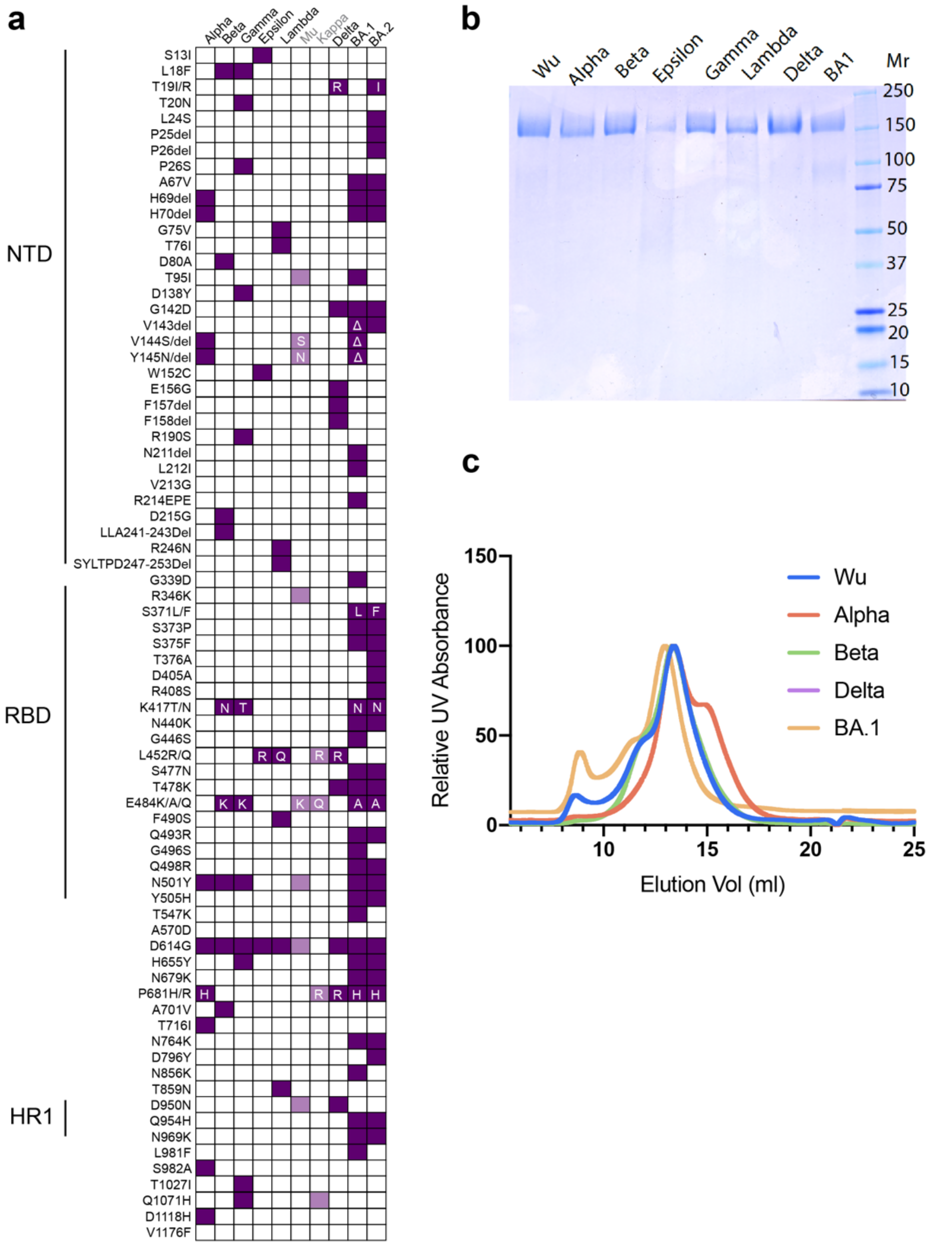


**Fig. S5. Mutations in VOC spike proteins and characterisation of the spike preparations**

**a** Mutations as labelled in purple were introduced in spike proteins. The black-colored spike variants are included in this study, the grey-colored ones not.

**b** The proteins were affinity purified, concentrated and buffer exchanged into PBS pH 7.4. Two micrograms of each protein were prepared in 4X lysis buffer, boiled and reduced. Proteins were loaded on SDS-PAGE and visualised using Coomassie Blue staining indicating pure purified spike protein.

**c** Size exclusion chromatography profiles of purified spike proteins (Superose 6 Increase 10/300 column).

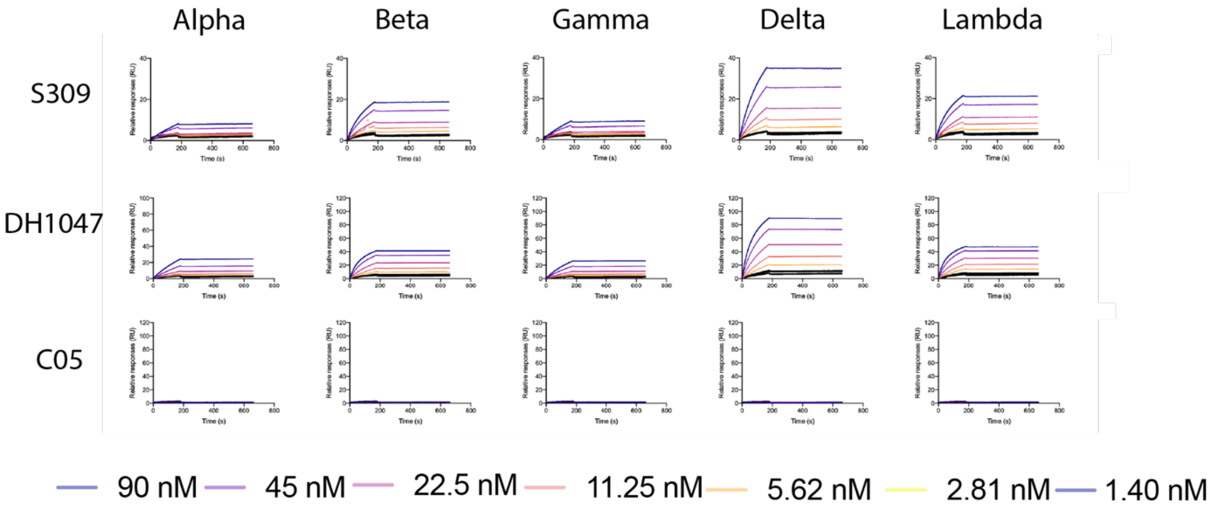


**Fig. S6. SPR sensorgrams of S309, DH1047 and C05**

mAbs were immobilized on SPR protein A chips. Various concentrations of SARS-CoV-2 spike variant proteins as indicated were injected for 180 s, followed by dissociation for 600 s. Dissociation constants ($K_D$) were determined on the basis of fits, applying a 1:1 interaction model. Dissociation constants ($K_D$) were determined based on fits, applying a 1:1 interaction model.

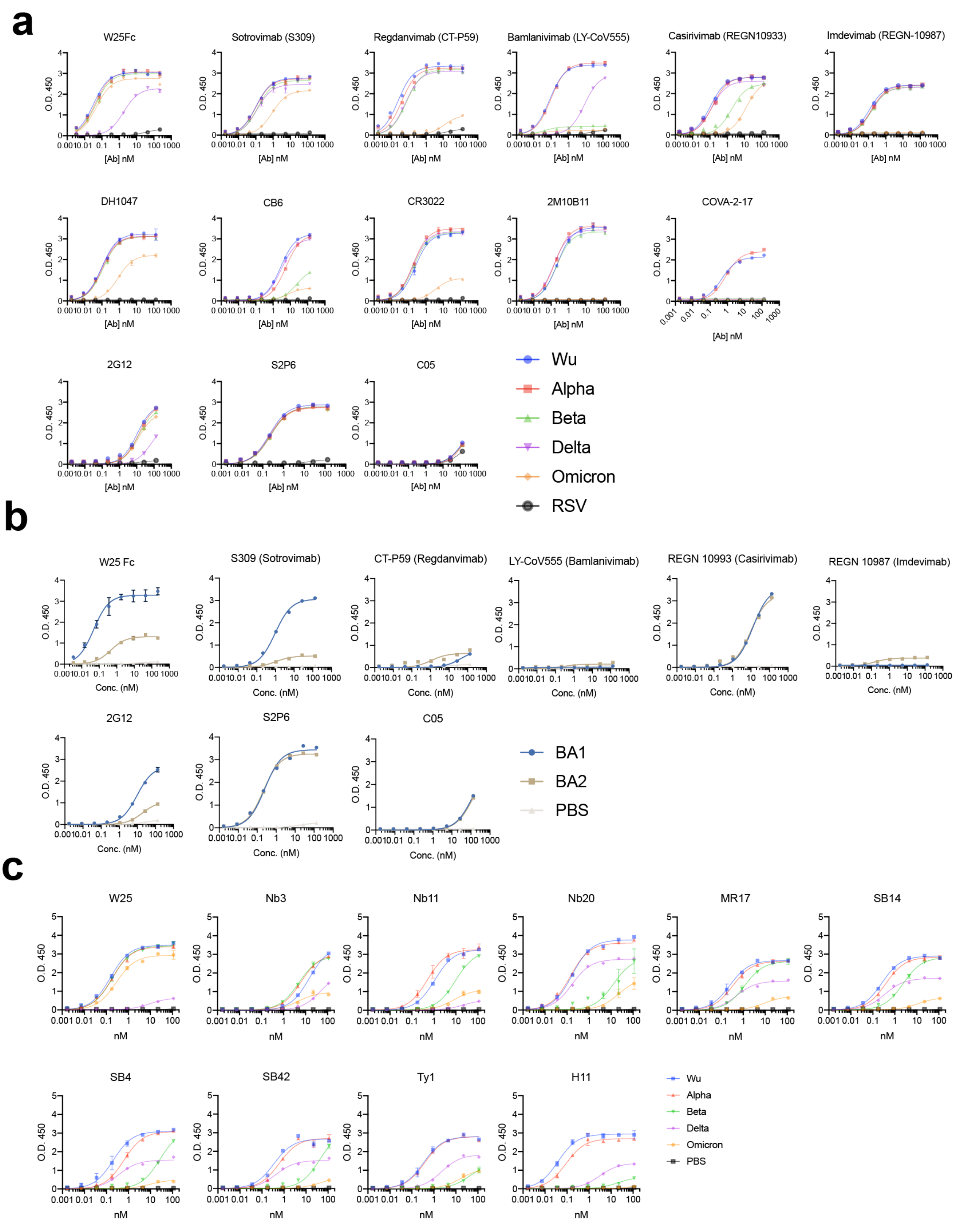


**Fig. S7. ELISA binding curves of W25Fc, EUA mAbs other epitope specific mAbs to SARS-CoV-2 Spikes variants**

**a** Binding curves of W25Fc, S309, CT-P59, REGN-10933, REGN-10987, DH1047, CB6, CR3022, 2M10B11, COVA2-17, 2G12, S2P6 and C05 (control mAbs) to Wu, Alpha, Beta, Delta and Omicron (BA.1).

**b** Binding curves of W25Fc, S309, CT-P59, REGN-10933, REGN-10987, 2G12, S2P6 and C05 (control mAbs) to BA.1 and BA.2**.**

**c** Binding curves of W25Fc, Nb3, Nb11, Nb20, MR17, SB14, SB4, SB42, Ty1, H11 to Wu-1, Alpha, Beta, Delta and Omicron (BA.1) demonstrating that W25 Fc binds Omicron spike whereas other Nbs lost their affinities to Omicron spike.

**a**

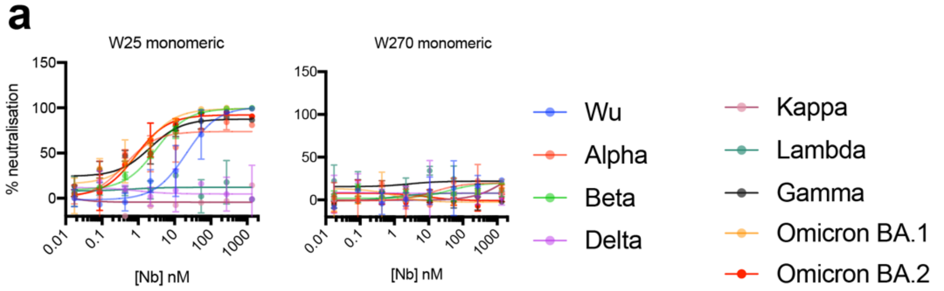


**b**

| Domain | IC$_{50}$ (nM±SEM) | | | | | | | | |
|---|---|---|---|---|---|---|---|---|---|
| | Wu | Alpha | Beta | Gamma | Kappa | Delta | Lambda | BA.1 | BA.2 |
| W25 | 22.63±7.97 | 0.42±0.22 | 2.10±0.35 | 2.12±0.98 | - | - | - | 0.62±0.22 | 0.85±0.26 |
| W270 | N/A | N/A | N/A | N/A | N/A | N/A | N/A | N/A | N/A |

**Fig. S8. Live virus neutralisation assay of W25 Nb against SARS-CoV-2 Wu and VOCs**
**a** Neutralisation curves comparing the sensitivity of SARS-CoV-2 strains (Alpha, Beta, Delta, Kappa, Lambda, Gamma) to W25 Nb, control Nb (W270) as indicated. Live SARS-CoV-2 virus different variants were incubated with serially diluted mAbs, in duplicate, for 1 hour at 37C, and then used to infect Vero E6 cells. After 30 mins, overlay media were added. Twenty hours post-infection, overlays were removed, cells were fixed with cold 80% acetone and dry prior staining for foci. The data were analysed and plotted using nonlinear regression (curve fit, three parameter) and IC$_{50}$ value was calculated from neutralisation curve by Graphpad Prism 8 software. Color represents SARS-CoV-2 variants.

**b** Summary of IC$_{50}$ values (nM) of neutralisation of SARS-CoV-2 variants performed in VeroE6 cells. Values that approached 50% neutralization were estimated from (A). Data are generated from two independent experiments, each performed in technical duplicate.

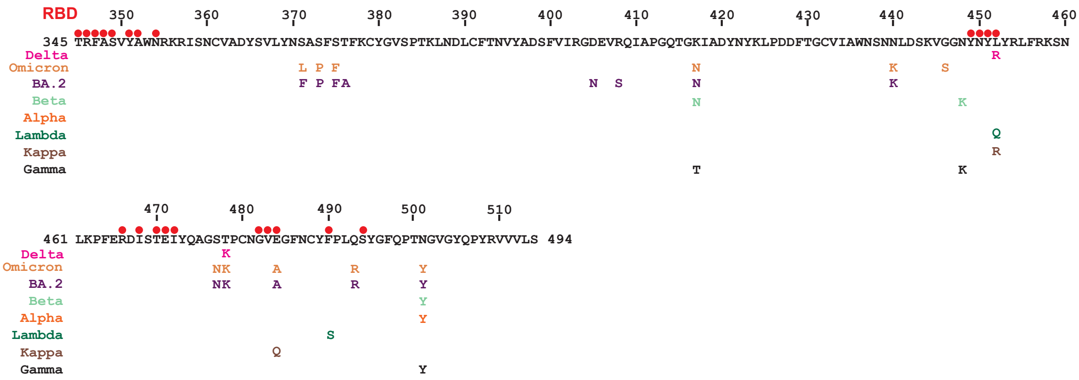

**Fig. S9. Amino acid residues on spike RBD involved in molecular contacts with W25**

Interacting residues are indicated as red dots above the sequence. Positions mutated in the SARS-CoV2 variants are shown in the sequence.

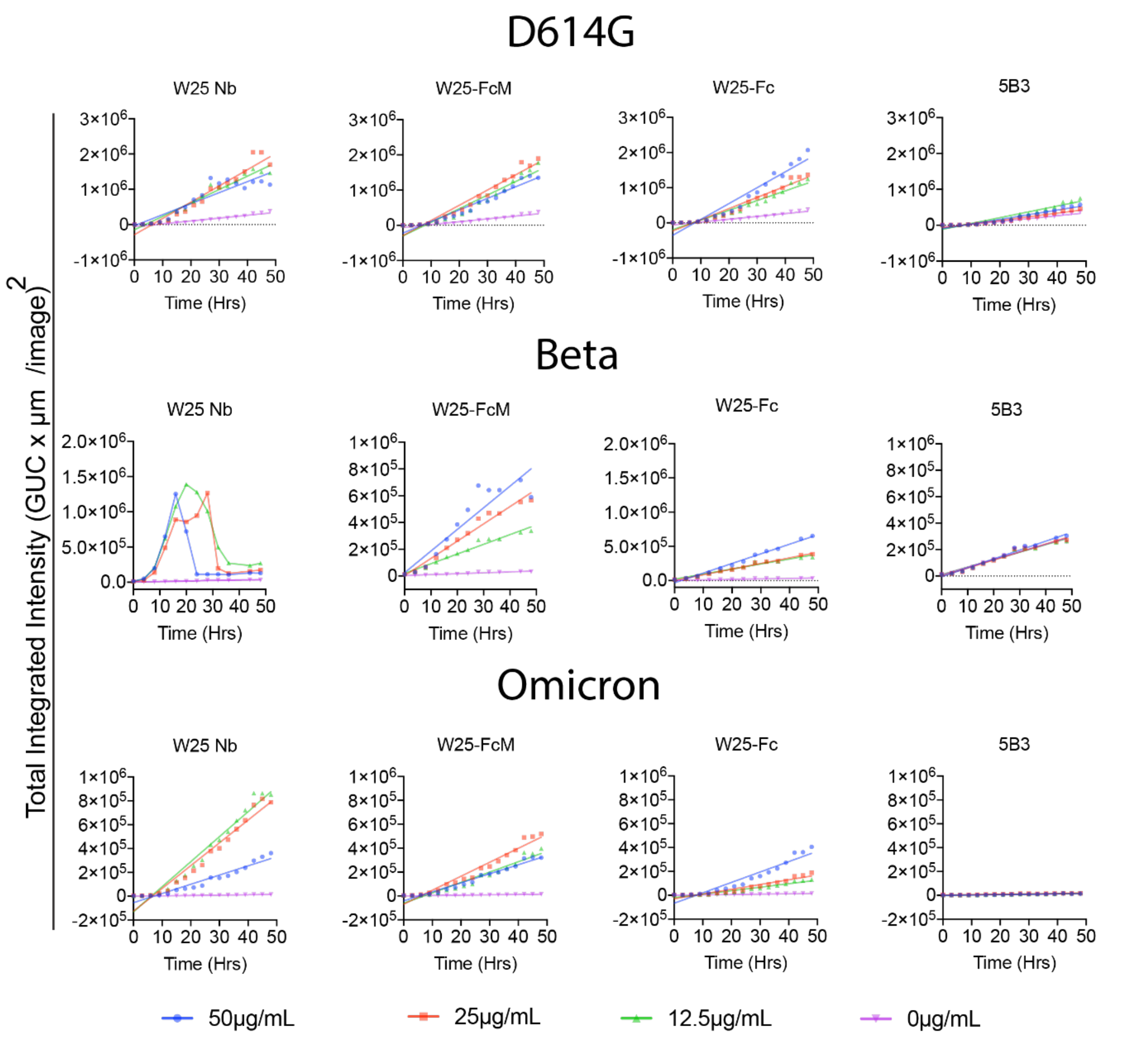


**Fig. S10. Cell-cell fusion assay of SARS-CoV-2 spike**

W25 enhances spike-mediated cell fusion monitored by GFP-positive syncytia cells. The data plotted shows the total sum of syncytia fluorescent intensity in the image, calculated used the total integrated intensity metric and expressed as green count units (GCU) per μm$^2$. W25, its derivatives and control antibody were diluted to the indicated concentrations and subsequently incubated with $2\times10^4$ effector cells in 50 μl at 37 °C, 5% $CO_2$ for 1 h. The W25 Nb, W25-FcM, W25-Fc, 5B3 (as control) and effector cell mixture were then subjected to incubation with target cells in corresponding wells and incubated for 18–24 h. GFP-positive syncytia were imaged every hour using the IncuCyte S3 live cell imaging system (Essen BioScience). Five fields of view were taken per well at 10× magnification, and GFP expression was determined using the total integrated intensity metric included in the IncuCyte S3 software (Essen BioScience). Representative images are illustrated in fig.S11.

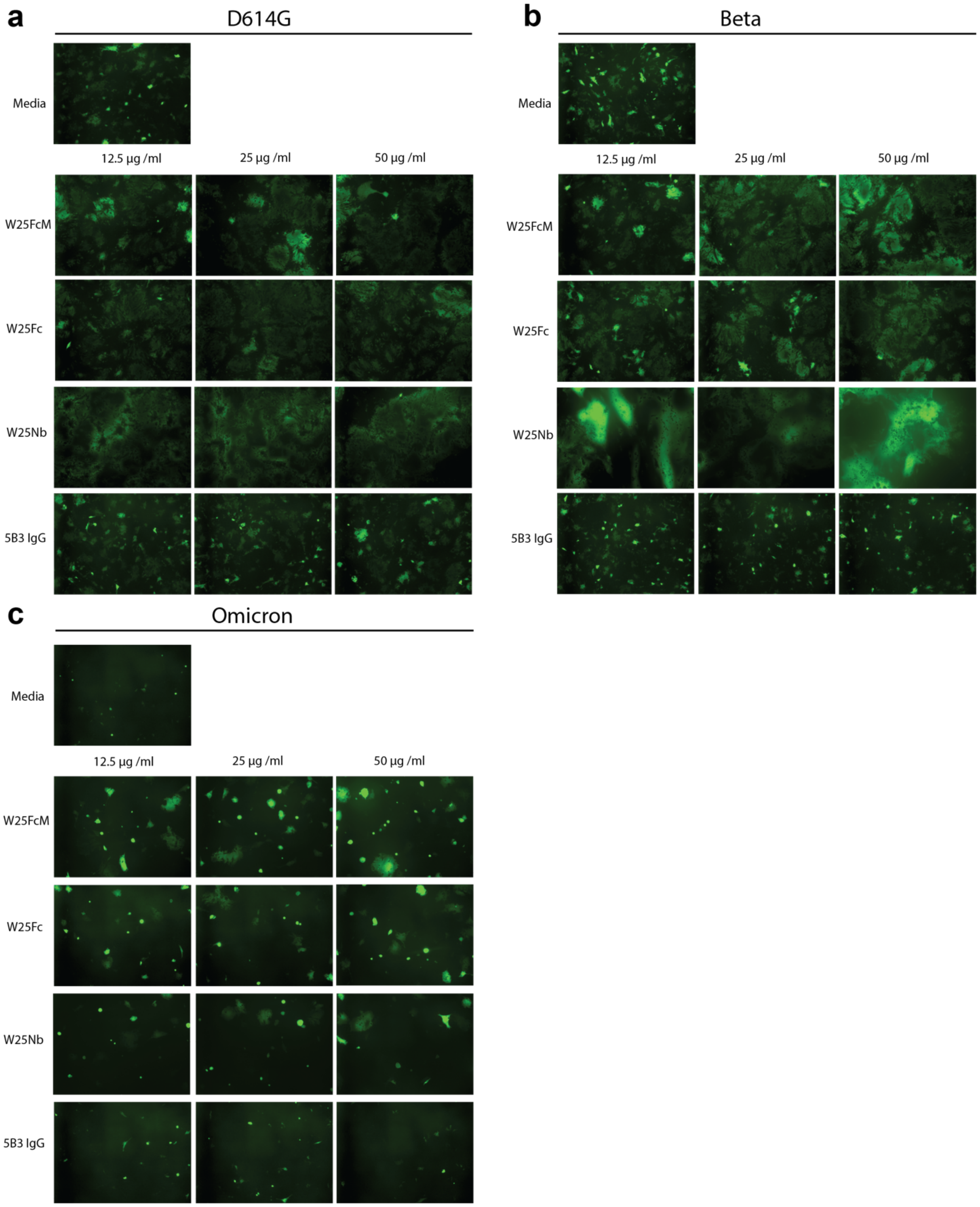


**Fig. S11. Representative images illustrating GFP-positive syncytia from cell-cell fusion corresponding to Fig. S10**

Cells were transfected with **a** SARS-CoV-2 D614G, **b** Beta and **b** Omicron variant spikes.

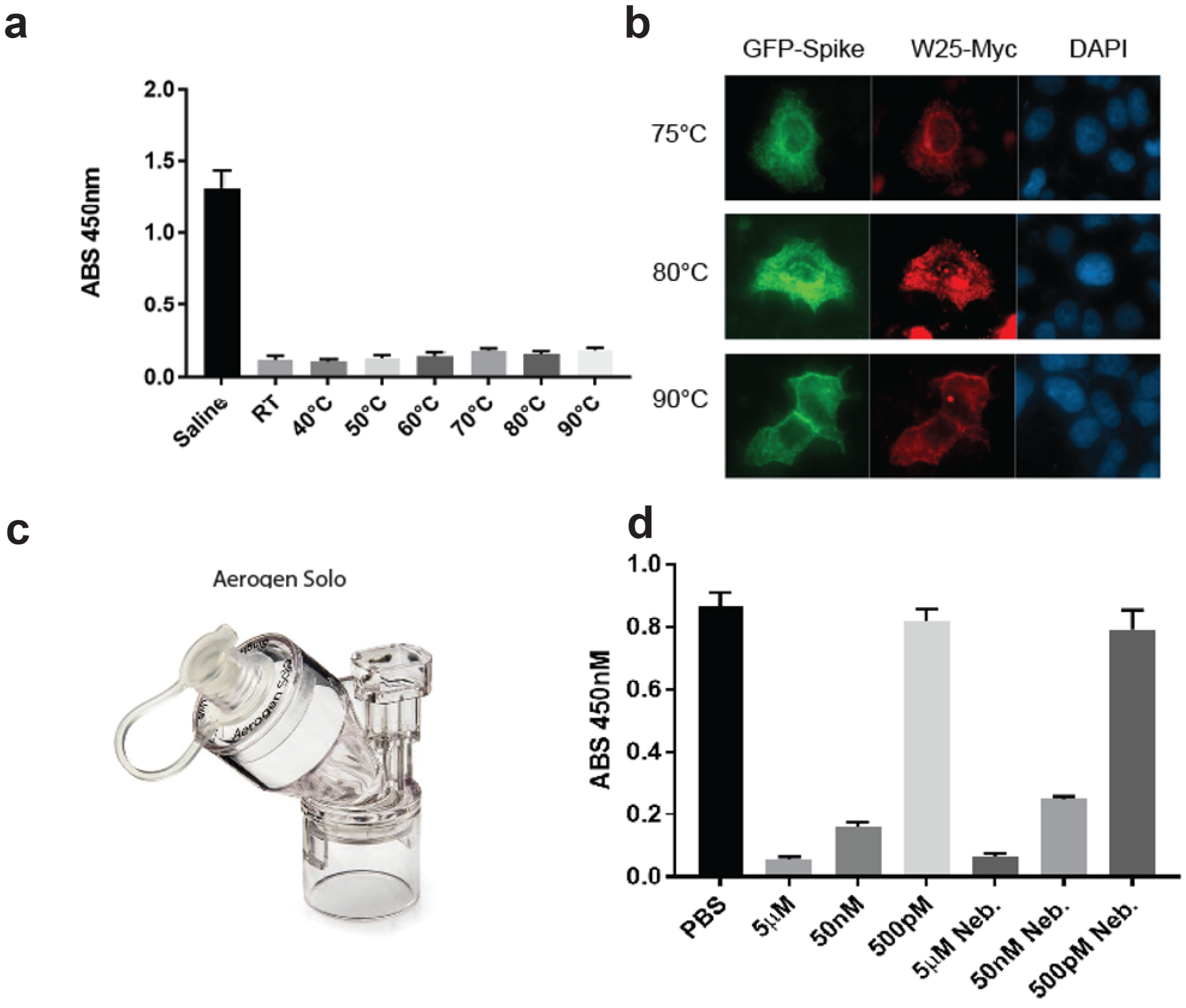


**Fig. S12. Thermostability and nebulisation stability of W25**

**a** Competition ELISA assay. RBD was immobilised on ELISA plates and W25, covalently modified with HRP, was added in the presence or absence of untagged W25, which was heat-treated for 20 min at the indicated temperatures.

**b** Immunofluorescence of Hela cells transient transfected with GFP-Spike using heat-treated myc tagged W25 (red).

**c** Aerogen Solo nebulizer used for W25 Nebulisation in PBS.

**d** Competition ELISA as in A, using decreasing amounts of W25 or W25 post-nebulisation

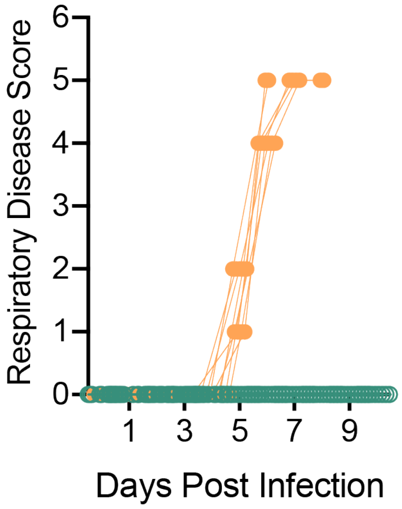


**Fig.S13 Respiratory disease scores of K18-ACE2 mice infected with SARS-CoV-2 Beta variant and received W25-Fc 4 h prior to infection**

Respiratory disease scores of SARS-CoV-2 Beta infected mice (Fig. 4A-E).

a

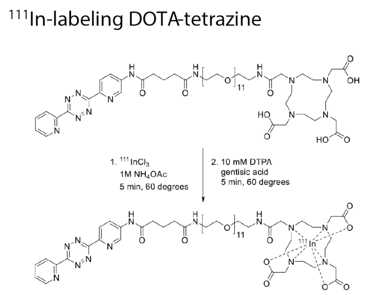


b

| | Time of euthanasia | Mice (n) |
|---|---|---|
| Group 1 | 5 min | 3 |
| Group 2 | 20 min | 4 |
| Group 3 | 1 hour | 4 |
| Group 4 | 3 hours | 4 |
| Group 5 | 5 hours | 4 |
| Group 6 | 24 hours | 4 |

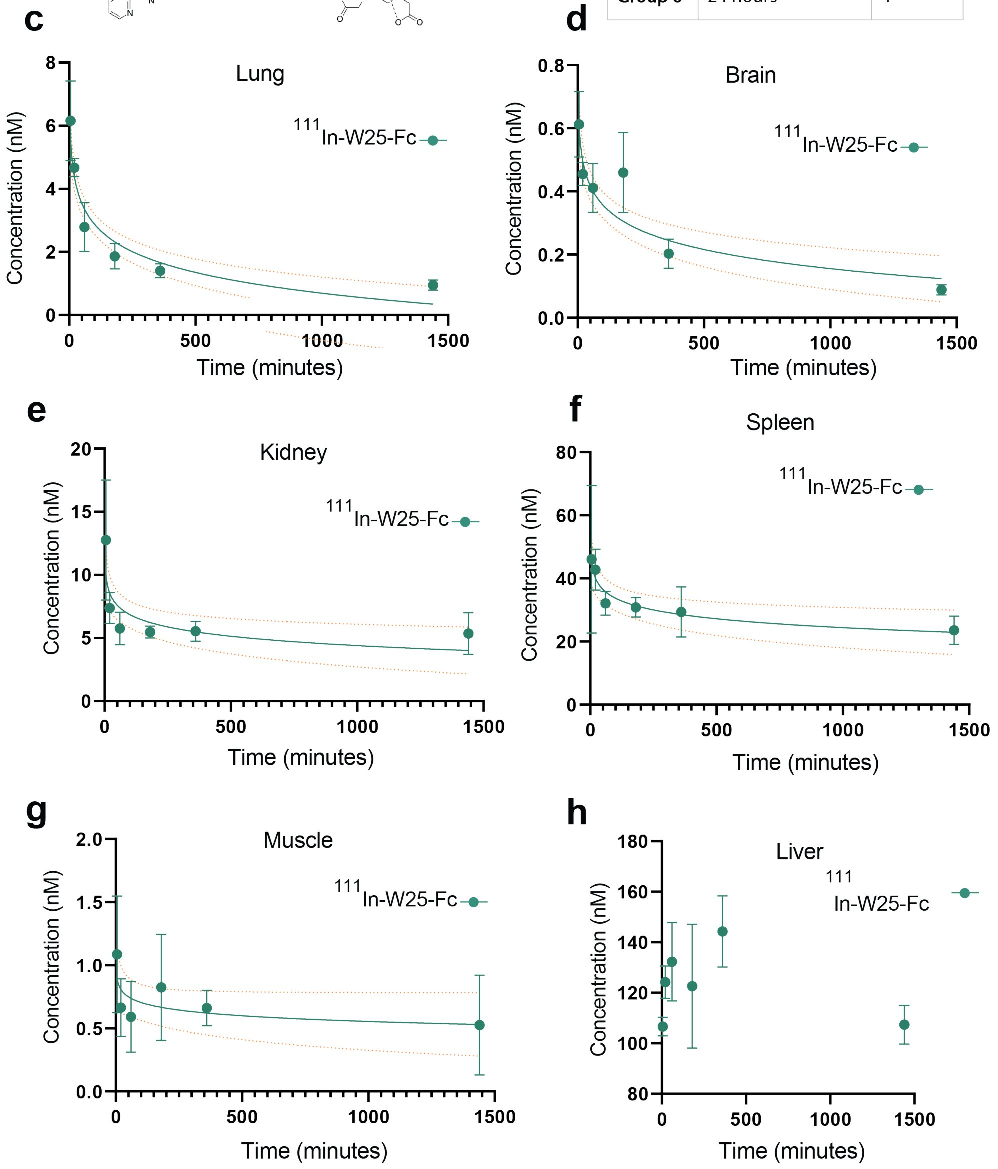


**Fig.S14. Pharmacokinetic of $^{111}$Indium radiolabelled W25-Fc**

**a** Conjugation of W25-Fc was conjugated to radioactive $^{111}$Indium.

**b** $^{111}$Indium W25-Fc (1 mg/kg) was injected intravenously via the tail to 6 groups of mice (group 1: 5 min, group 2: 20 min, group 3: 60 min, group 4: 3 hr, group 5: 5 hr and group 6: 24 hr). The mice were dissected and concentration of $^{111}$In W25 was measured the following tissues using an Auto-Gamma Counter: **c** Lung, **d** Brain, **e** Kidney, **f** Spleen, **g** Muscle and **h** Liver.

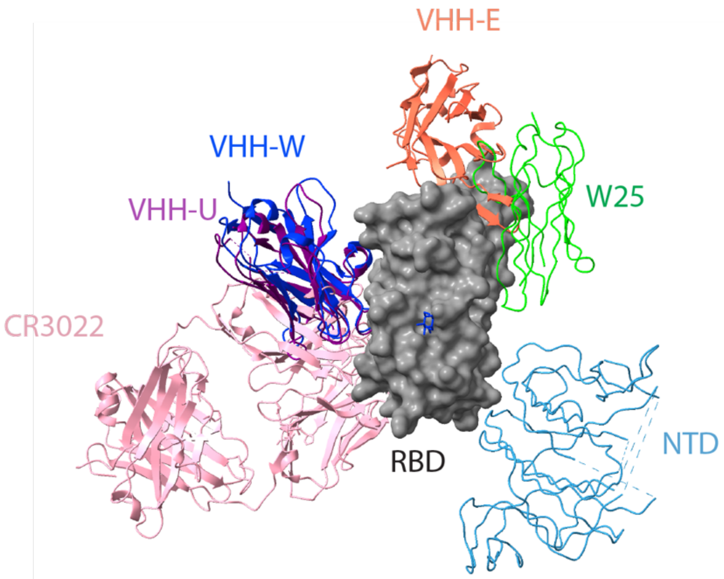


**Fig. S15. Structural comparison of antibodies and nanobodies that induce unconventional neutralisation mechanism**
W25, VHH W (PDB 7KN7), U (7KN5), and E (7KN5), and mediating spike disruption: CR3022 (6W41).

**Supplementary Movie S1.** The movie illustrates the cryo-EM structure of the Wu spike/W25 complex, including a close-up of the RBD/W25 molecular model, highlighting the interaction interface.
